# A deep population stratification and ongoing local adaptations of the range-expanding lovebug *Plecia longiforceps*

**DOI:** 10.1101/2025.07.02.662813

**Authors:** Donghee Kim, Jonghwan Choi, Dongyoung Kim, Marie-Laurence Cossette, Sangil Kim, Cheng-Lung Tsai, Min Jeong Baek, Sun-Jae Park, Amaël Borzée, Seunggwan Shin, Choongwon Jeong

## Abstract

Rapidly range-expanding species provide a superb opportunity to study dynamic demography and ongoing adaptations to novel environments. *Plecia longiforceps*, a species of lovebug flies native to southeast China and Taiwan, has recently been expanding in Okinawa in Japan and Seoul in South Korea, where massive outbreaks are garnering public attention. Here we analyzed whole-genome sequences of 150 individuals across China, Korea, Okinawa, and Taiwan, including a newly identified northern Chinese population from Qingdao. We found a deep divergence between continental (Korea-China) and insular (Okinawa-Taiwan) populations and specify northern China and Taiwan as sources for Korea and Okinawa, respectively. Severely reduced genetic diversity in Korea and Qingdao suggests serial founder events during their northward expansion. An old split time between Qingdao and Korea (∼3,500 generations ago) and multiple gene flows into Qingdao reject a simple scenario of a single recent northward migration. Selection signals in northern groups include genes associated with pigmentation, thermal sensing, lipid metabolism, and immunity. We observed a markedly reduced genetic diversity on X chromosome in northern populations, indicating ongoing strong sex-biased selection. Our findings portrait details of the northward range expansion in the species and highlight the role of adaptation for successful invasion at higher latitudes.

## Introduction

Range-expanding organisms serve as important indicators of the anthropogenic impacts on biodiversity. Warmer climates drive range shifts to higher altitudes and latitudes^1–3^, while human activities move species into non-native habitats^4,5^. Newly introduced species interact with the local ecosystem, sometimes causing devastating effects^6,7^. Furthermore, they can lead to economic challenges by causing extensive damage to agriculture, threatening public health, and/or becoming public nuisances^8–10^. Therefore, when a non-indigenous species begins to proliferate far from its native range, identifying the origin of this population and tracing its route of introduction becomes a priority to evaluate its ecological and socioeconomic impacts^11–13^.

Species dispersing into new habitats are confronted with various environmental challenges, and therefore have gained scientific interest as models for exploring rapid evolutionary processes^14–16^. Populations at the expanding front tend to have lower genetic diversity due to founder effects^13^, best represented in long-distance colonization where small founding populations experience severe bottlenecks^17,18^. The ability of such populations to overcome genetic constraints and thrive in novel environments has been widely debated, a phenomenon known as the “genetic paradox of invasion”^19,20^. Several mechanisms may explain invasion success despite initial bottlenecks: i) repeated introductions from the same or different source populations promotes admixture, restoring genetic diversity^12^; ii) hybridization with native species introduces novel and adaptive alleles^21^; iii) strong selection on existing or novel mutations enables rapid adaptation despite low genetic variation^22^; and iv) in rare cases, genetic bottlenecks may increase additive genetic variance by disrupting co-adapted gene complexes^23^. The ongoing expansion of the lovebug *Plecia longiforceps* in East Asia provides a well-defined case to study the rapid evolution of a range-expanding species in action. Indeed, the genus *Plecia* includes other well-documented cases of range expansion, such as *P. nearctica* in North America, offering a valuable comparative system^24^. Originally known from the subtropical woodlands of southeast China and Taiwan, *P. longiforceps* has been sequentially colonizing the Ryukyu Islands over the last several decades^25–27^. Since 2022, massive outbreaks have made this insect a serious nuisance pest in the Seoul Metropolitan Area of South Korea^28^. The annually recurring and intensifying swarms of *P. longiforceps* in Korea present not only public disturbances but also environmental concerns, as population explosions can indicate ecological disequilibrium after biological invasions^29,30^, while also impacting resource availability and allocation among local species^31,32^. Previous studies on *P. longiforceps* have investigated potential future range shifts based on climate change models^28^, microbial composition as possible public health risks^33^, and genomic content with implications on urban habituation^34^. However, it requires population-scale data collected throughout its native and introduced ranges to uncover the patterns of dispersal and establishment into novel environments.

The environmental differences across its expanding distribution suggest the possibility of adaptive evolution in *P. longiforceps*. With a small body (∼7 mm) and short adult lifespan (∼1 week), this insect has overcome numerous obstacles between its native range and newfound settlements, including the transition from subtropical to temperate climates, dispersal over the Yellow Sea, and acclimation to Seoul’s urban habitat. In particular, climatic shifts are expected to be an important driving force for local adaptation in *P. longiforceps*. Both temperature and thermal susceptibility are major factors in determining an insect’s geographical range^35,36^, and genetic signatures of climatic adaptation, such as cold tolerance, have been repeatedly identified in species expanding into higher latitudes^22,37–39^. Meanwhile, *P. longiforceps* exhibits different flight seasons by latitude. Adults have been recorded throughout the year in Taiwan, whereas in southern China and Okinawa biannual emergence is reported in spring and fall, suggesting seasonal influence^25–27^. In Korea, outbreaks only occur once in summer^28^. Such rapid phenotypic changes are often assumed to be a case of innate developmental plasticity that allows insects to withstand environmental variations, but can also arise due to rapid genetic changes by natural selection^22^. Thus, population-scale genome variation data can provide fundamental resources for interpreting adaptive evolutionary processes.

In this study, we aim to unravel the range expansion process of *P. longiforceps*, focusing on its demographic history and adaptive evolution. We present whole-genome resequencing data for 150 specimens to represent four regions from its native range in southeast China (Nanjing and Hangzhou) and Taiwan (Nantou and Pingtung), a newly discovered population in eastern China (Qingdao), as well as two recently introduced localities: Japan (Okinawa) and Korea. Utilizing over 16 million single nucleotide polymorphisms (SNPs), we provide a fine-grained reconstruction of the genetic history of *P. longiforceps* populations regarding their dispersal and gene flow, and detect clear signatures of adaptive evolution in many genes relevant to high-latitude adaptations.

## Results

### Population-scale genome variation data from *P. longiforceps*

We obtained whole genome resequencing data from 156 *P. longiforceps* individuals collected across 32 sampling sites in seven major regions (Fig. 1). Reads were aligned to the Korean *P. longiforceps* reference genome^34^, resulting in an average sequencing depth of 13.6x (Table S1). After filtering out 6 individuals with low endogenous DNA (< 40%) or high bacterial contamination (> 25%), we called variants from 150 individuals, identifying 44,877,370 biallelic single-nucleotide polymorphisms (SNPs) and 6,929,490 short insertions/deletions (≦ 40 base pairs).

**Fig. 1.**
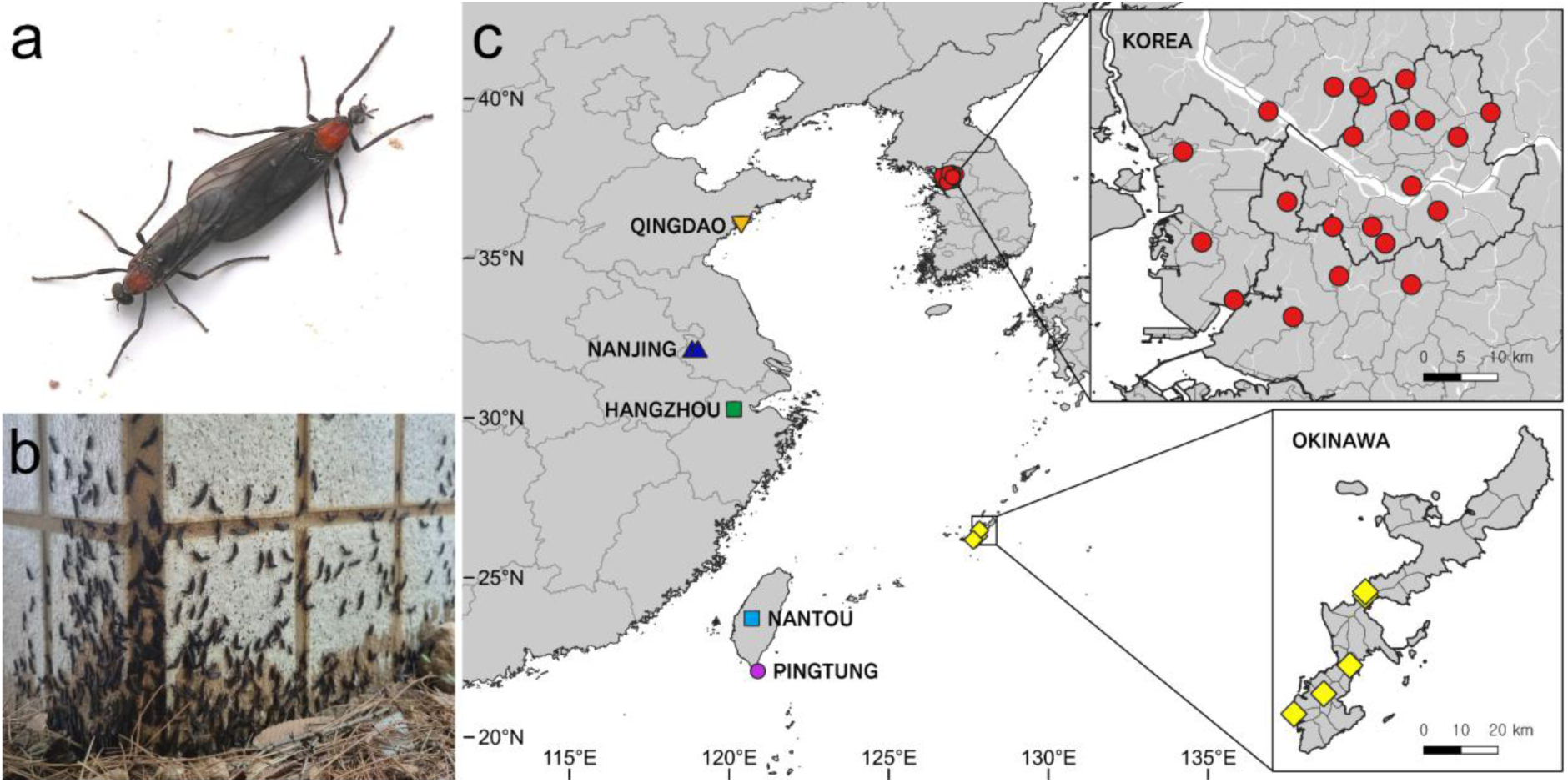
Population sampling of *P. longiforceps*. **a**, A pair of *P. longiforceps* in copulation. **b**, Mass emergence of adult lovebugs in Seoul, Korea, observed on June 19th, 2024. **c**, Sampled locations of *P. longiforceps* in East Asia. Detailed views for Korea (upper right) and Okinawa (lower right) are provided as insets. Marker shapes and colors correspond to Fig. 2a. Maps are sourced from GADM version 2.5 (https://gadm.org/) and the Republic of Korea Ministry of Land, Infrastructure and Transport.

The proportion of the genome covered when mapped to the reference varied among populations, hinting at highly diverged genetic profiles between the regions. Korean individuals showed the highest median genome coverage (84.8%), followed by Qingdao (84.4%), Nanjing (80.2%), Hangzhou (79.3%) in China, Nantou in Taiwan (67.3%), and Okinawa in Japan (64.1%) (Fig. S1, Table S1). Samples from Pingtung in Taiwan (n=5), sourced from museum specimens, showed the lowest coverage (median = 14.9%) and unusually low sequencing depth (median = 1.7x), due to poor preservation of DNA.

To assess variation in mappability across populations, we identified population-specific low-mappability regions where sequencing depth was below one fifth of the genome-wide average. Overall, these regions were enriched in repetitive and GC-biased sequences (Figs. S1, S2, and S3). In the Okinawa population, for example, the repeat content was significantly higher in low-mappability regions compared to mappable regions (median=0.91 vs. 0.60, p<2.2×10^-16^, Welch’s t-test). Additionally, both high- and low-GC regions were more likely to be unmappable, as indicated by the GC content’s standard deviation being twice as high in low-mappability regions compared to mappable ones (0.10 vs. 0.05, p<2.2×10^−16^, Brown-Forsythe test).

**Fig. 2.**
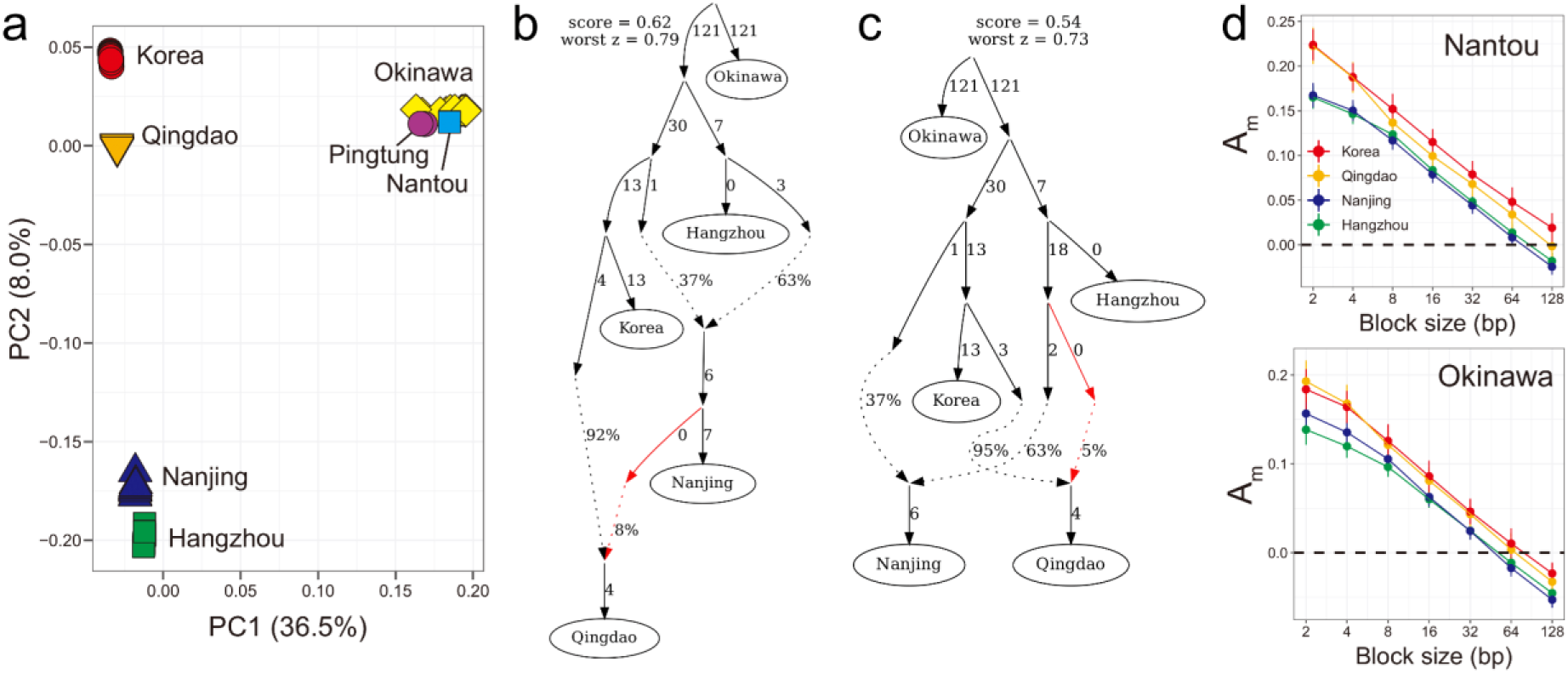
Population structure in *P. longiforceps*. **a**, Principal component analysis (PCA) of 150 East Asian *P. longiforceps* diploid genotypes. Each individual is represented by a color-filled symbol corresponding to its group. The proportion of variance explained by each principal component is indicated on the axis labels. **b–c**, Representative population graphs inferred using qpGraph, showing shared southern ancestry between Qingdao and Nanjing. In **b,** Nanjing directly contributes to Qingdao. In **c,** the southern ancestry of Qingdao is closer to that of Nanjing compared to Hangzhou. In both panels, the southern lineage contributing to Qingdao is marked as red. Edge lengths correspond to *F_ST_* × 1000. The graph’s log-likelihood score and the largest residual deviation between observed and expected *f*-statistics (“worst *z*”) are displayed above the graph. **d,** Estimates of A_m_ for various block sizes between island (recipient) and continent (donor) populations. We used a single high-coverage Nantou genome (upper) and Okinawa genome (lower) as representatives of island populations. Colored points and lines correspond to the continental population used to estimate A_m_. Error bars correspond to the 95% CIs of each estimate.

To reduce noise and errors in our genotype data set, we filtered out SNPs in regions with low-mappability, high repeat content, or extreme GC content, and applied additional quality control measures and retained 16,989,044 high-quality SNPs for further analyses (see Methods for details).

### A deep population structure of *P. longiforceps*

To explore the genetic diversity of East Asian *P. longiforceps* populations, we performed a principal component analysis (PCA) using diploid genotypes from 150 individuals across 16,989,044 SNPs. PC1 separates the Korea-China populations from Taiwan-Okinawa while PC2 delineates the northern (Korea and Qingdao) and southern (Hangzhou and Nanjing) groups within the Korea-China populations (Fig. 2a). PC1 and PC2 explained 36.5% and 8.0% of the total variation, respectively. Within the Korea-China populations, the distribution of Korea, Qingdao, Nanjing, and Hangzhou populations along PC2 correlates with their geographic latitudes, from north to south (Fig. 1, Table S1). These isolation-by-distance patterns are further supported by genetic differentiation, as measured by pairwise *F*_ST_ values (Korea-Qingdao *F_ST_* = 0.16; Korea-Nanjing *F_ST_* = 0.30; Korea-Hangzhou *F_ST_* = 0.33) (Table S2).

To infer the phylogenetic relationship between the *P. longiforceps* populations, we tested for evidence of differential gene flow between the northern and southern populations using *f*_4_-statistics in the form of *f*_4_(Okinawa, pop1; pop2, pop3). First, *f*_4_(Okinawa, Korea/Qingdao; Hangzhou, Nanjing) ≧ 53.1 standard error measures (s.e.m.) show that the northern populations are genetically closer to Nanjing than to Hangzhou, mirroring geographic distance (Table S3). Second, *f*_4_(Okinawa, Hangzhou; Korea, Qingdao) = 5.8 s.e.m. shows that Korea and Qingdao do not form a clade and that there is gene flow between the geographically closer Qingdao and Hangzhou (Table S3).

QpGraph^40^ modeling supports two topologies where both involve a substantial contribution (∼40%) from the northern group into the Nanjing population: (1) Qingdao receives ∼8% gene flow directly from the admixed Nanjing population (Fig. 2b), or (2) Qingdao inherits ∼5% of its ancestry from the southern ancestral source of Nanjing (Fig. 2c). Further supporting Nanjing as the most likely southern source, Momi2 analysis demonstrated that a model with gene flow from Nanjing into Qingdao provided a better fit than one involving Hangzhou (Fig. 4b, Table S5). Additionally, introgressed variants in Qingdao that are absent in Korea are more frequently shared with Nanjing than with Hangzhou individuals (Fig. S7). Alternative topologies without shared southern ancestry between Qingdao and Nanjing show significantly worse fits (worst |*f*_4_| > 4.2 s.e.m) (Fig. S4).

The PCA confirms a close match between the genetic profiles of Okinawa and Taiwan, thus implies a Taiwanese origin of the Okinawan population. Interestingly, our sample set, although limited to only two locations in Taiwan, provides information on the fine-scaled geographic origin of the Okinawan population from Taiwan. First, Okinawa is genetically homogenous as shown by |*f*_4_(Korea, Nantou/Pingtung; Okinawa 1, Okinawa 2)| ≦ 2.5 s.e.m. for 11 of 12 comparisons, consistent with a discrete clustering pattern observed in PCA (Table S3). Second, Okinawa is closer to Nantou than to Pingtung as shown by *f*_4_(Korea, Okinawa; Pingtung, Nantou) = 22.9 s.e.m. (Table S3). However, a simple tree-like relationship of (Pingtung, (Nantou, Okinawa)) is also rejected. *F*_4_-statistics reveal that Okinawa is genetically intermediate between Pingtung and Nantou: *f*_4_(Korea, Nantou; Pingtung, Okinawa) = 46.6 s.e.m. and *f*_4_(Korea, Pingtung; Nantou, Okinawa) = 9.4 s.e.m. (Table S3). Therefore, a single origin from an unsampled Taiwanese population whose genetic profile is intermediate between Nantou and Pingtung is a parsimonious scenario for the origin of the Okinawan populations.

The PCA also reveals strong genetic differentiation between continental (Korea-China) and island (Okinawa-Taiwan) populations, reflected by limited shared polymorphism (only 4.8% of total variants) and high pairwise *F*_ST_ values (e.g., Korea-Okinawa *F*_ST_ = 0.73) (Fig. 2a, Tables S2, S4). Despite this differentiation, we found evidence of gene flow between these groups. First, *f*_4_(Hangzhou/Nanjing, Qingdao/Korea; Pingtung, Nantou/Okinawa) significantly fall below zero (*f*_4_ ≦ - 20.5 s.e.m.), indicating gene flow between at least one of the following pairs: (1) Hangzhou/Nanjing and Nantou/Okinawa, or (2) Qingdao/Korea and Pingtung (Table S3). Second, we inferred the direction of gene flow between continental and island populations using the two-taxon A_m_ statistic (Mackintosh and Setter 2024). The A_m_ statistic measures scaled heterozygosity differences between individuals from different populations, leveraging the fact that gene flow increases heterozygosity in the recipient while remaining unchanged in the donor. To account for population size effects, A_m_ is calculated in genomic blocks where both individuals are heterozygous, indicating a conflicting topology caused by either (1) incomplete lineage sorting or (2) gene flow. In the first case, both individuals have equal heterozygosity. In the second, the recipient retains higher heterozygosity, making the A_m_ value positive. We restricted A_m_ calculations to blocks size ≦ 128 bp to minimize recombination effects. For all continental-island pairs, A_m_ values were significantly positive (recipient: island) at short block sizes (≦ 32 bp), supporting gene flow from the continent to the island (Fig. 2d). As block size increased, A_m_ values declined due to recombination disrupting linkage.

### Strong serial bottleneck during the northward range expansion

Populations that have recently expanded their range through a series of founder events are expected to exhibit a gradual decrease in genetic diversity along the direction of expansion^41^. In accordance with the hypothesis of a south-to-north expansion, the Korea-China *P. longiforceps* populations (Korea, Qingdao, Nanjing, and Hangzhou) show a strong negative correlation between genome-wide heterozygosity, a measure of genetic diversity, and latitude: Korea (4.0±0.4 sites per 1,000 base pairs (bp), mean ± 1σ), Qingdao (5.7±0.2), Nanjing (8.3±0.3), and Hangzhou (9.3±0.3) (Fig. 3a). In the Okinawa-Taiwan *P. longiforceps* populations, the Okinawa population (6.1±0.5) shows lower heterozygosity than the single Nantou genome (8.0±0.2), which may reflect a bottleneck during the dispersal into Okinawa from Taiwan.

**Fig. 3.**
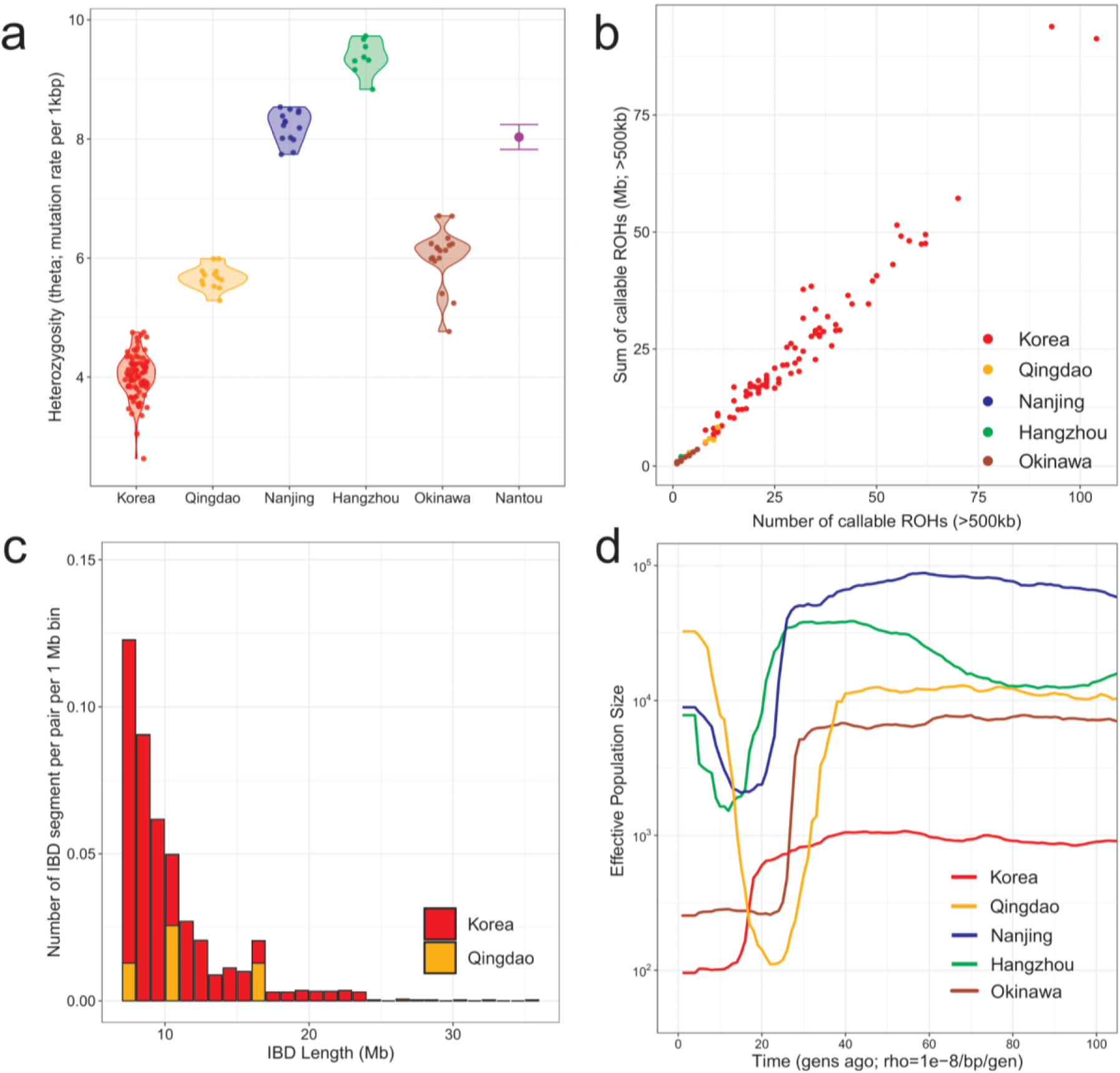
Strong bottleneck during invasion into Korea and Qingdao. **a**, Genome-wide heterozygosity (y-axis) per individual, calculated at the base-pair level from the callable genome, across populations (x-axis). Each colored point represents an individual. For the single Nantou genome, block jackknife error bars corresponding to the 95% confidence intervals are displayed. **b**, The total length of the callable genome in runs of homozygosity (ROH) ≥ 500 kbp (y-axis) plotted against the number of callable ROH blocks per individual (x-axis). **c**, Number of identity-by-descent (IBD) segments per pair (y-axis) longer than 7 Mb, plotted along IBD segment length. **d**, Effective population size (Nₑ) over the past 100 generations, reconstructed using GONE. The curve shows historical Nₑ (y-axis) over time (x-axis). In **a**–**d**, Colors indicate different population groups.

Likewise, Korea, Qingdao, and Okinawa individuals harbor long runs of homozygosity (ROH) blocks exceeding 500 kb, a hallmark of small effective population size. This effect is most pronounced in Korea: Korea (30.6 ROH blocks per individual; n=83 individuals with coverage > 8x), Qingdao (5.1 ROH blocks per individual; n=13), and Okinawa (1.9 ROH blocks per individual; n=17). In contrast, individuals from Hangzhou (n=8) and Nanjing (n=14) exhibited very few ROH blocks, with an average of 0.25 and 0.14 ROH blocks per individual, respectively. The total ROH block length per individual in the Korean population (25.0±16.2 Mb, mean ± 1σ) was likewise 8-fold greater than that in Qingdao (3.3±2.3 Mb). Additionally, we identified long identity-by-descent (IBD) segments > 7 Mb between individuals using IBIS^42^, with substantial IBD sharing within the Korean and Qingdao populations but none shared between populations or within other populations (Fig. 3c). The number of IBD segments per pair in Korea (n=0.40 per pair, average length = 10.3 Mb; total pairs = 3,403) was 8-fold greater than that in Qingdao (n=0.05 per pair, average length = 11.1 Mb; total pairs = 78). These patterns of extensive ROH within individuals and elevated IBD sharing between individuals strongly support a recent and severe bottleneck in both Korea and Qingdao, with the bottleneck in Korea appearing much more extreme.

To further explore the timing and magnitude of these bottlenecks, we estimated the effective population size (N_e_) over the last 100 generations for each population using GONE^43^, based on genome-wide linkage disequilibrium (LD) and assuming a recombination rate of 1 cM/Mb. The analysis revealed extreme declines in N_e_ to approximately 100 individuals in Korea (10 generations ago) and Qingdao (25 generations ago), and 250 individuals in Okinawa (25 generations ago) (Fig. 3d). In contrast, the Nanjing and Hangzhou populations experienced milder bottlenecks (ca. 2,000 individuals, 15 generations ago). Notably, the Qingdao population had a significantly larger N_e_ than Korea (10,000 vs. 1,000) prior to the bottleneck, suggesting distinct demographic histories between these northern populations. Since LD decays more rapidly at higher recombination rates, our use of conservative rate (1 cM/Mb vs 2.06–3.44 cM/Mb in *Drosophila* spp.) likely overestimates recent N_e,_ implying that the actual bottlenecks may have been even more severe^44^.

### The origins and demography of the northern populations

To investigate the long-term demographic history of *P. longiforceps*, we reconstructed the N_e_ over time using MSMC2^45^. We analyzed two high-coverage genomes per population (>19x; >13x for Okinawa), except for Nantou, where only one genome was available. We applied the mutation rate of *Drosophila melanogaster* (5×10^-9^ mutations per bp per generation)^44^.

In the Korea-China populations, all groups exhibited similar demographic trajectories, characterized by gradual population growth until approximately 100,000–200,000 generations ago, followed by a steady decline. However, the northern populations (Korea and Qingdao) experienced a rapid decline compared to the southern populations (Nanjing and Hangzhou) (Fig. 4a). Since approximately 40,000 generations ago, northern populations have maintained a small N_e_ of fewer than 50,000, consistent with their low genetic diversity.

**Fig. 4.**
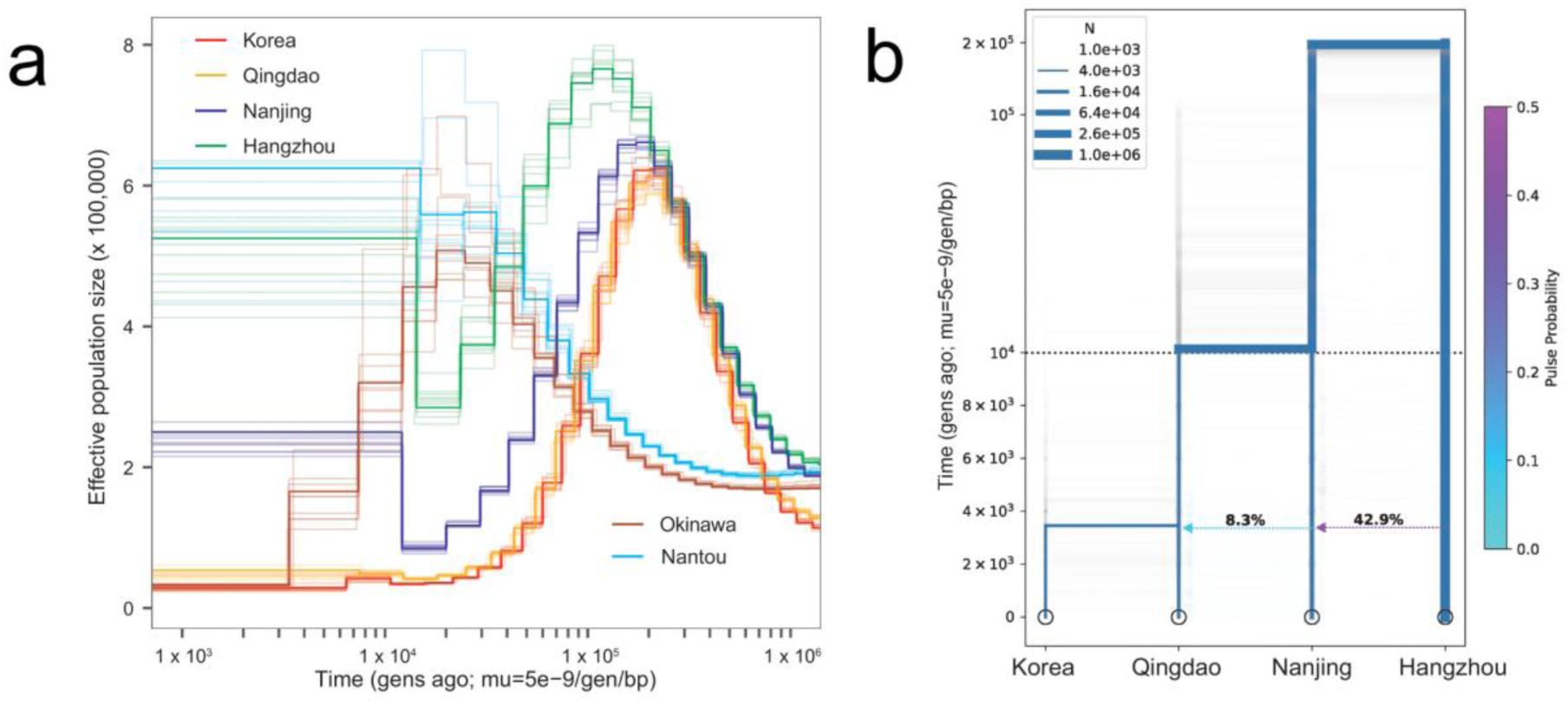
Demography and population size history. **a**, Long-term effective population size (*Nₑ*) from 1,000 to 1,000,000 generations ago, reconstructed using MSMC2. Each colored curve represents the demographic history of an *P. longiforceps* population (*y*-axis) over time (*x*-axis). Bootstrapped estimates are shown as transparent lines, while main runs are plotted as solid lines. A mutation rate of 5.0 × 10⁻⁹ per generation per base pair was assumed. **b**, Best-fit demographic model inferred using Momi2, depicting a gene flow from Nanjing into Qingdao. This model had the highest log-likelihood among all tested two-pulse scenarios. Maximum likelihood (ML) estimates are shown.

The Okinawa-Taiwan populations showed continuous growth until around 10,000-20,000 generations ago. However, while the Nantou population remained stable, the Okinawa population underwent a sharp population decline, reducing its N_e_ from around 500,000 to 30,000.

To estimate divergence times within the Korea-China populations, we applied the coalescent-based method Momi2^46^, using the Okinawa population as an outgroup to polarize ancestral alleles. Initially, we modeled Nanjing as a hybrid between the northern lineage and Hangzhou, while Qingdao remained cladal with Korea. This base model (Model 0) estimated that 38.9% of Nanjing’s ancestry derived from Hangzhou and placed the Qingdao-Korea divergence at ca. 9,000 generations ago (Fig. S5, Table S5), largely consistent with pseudodiploid X chromosome analysis using MSMC2 (ca. 8,000 generations ago) (Fig. S6). We then tested three alternative models incorporating gene flow into Qingdao from Nanjing (Model 1), Hangzhou (Model 2), or a basal lineage (Model 3) diverging from the ancestral population of all Korea-China populations. Each model provided a significantly better fit than the base model (p < 2.2 × 10^−16^, likelihood ratio test) and produced more recent Qingdao-Korea divergence estimates (Figs. 4b, S5, Table S5).

Among these, the best-fitting model (Model 1; AIC difference with Model 2 and 3 ≧ 1449.8) identified Nanjing as the source of southern gene flow into Qingdao, with an estimated 8.3% contribution (Fig. 4b). However, even in the model taking into account this southern gene flow, Qingdao and Korea remain substantially diverged by ca. 3,500 generations (95% CI: 889–10,914), suggesting that i) the sampled Qingdao population does not represent the true source of the Korean population and ii) northern populations do not stem from a recent expansion of a single ancestral source.

Given the limited geographic sampling of southern populations, we further tested whether an unsampled basal lineage contributed to Qingdao ancestry. The three-way admixture model (Model 4), which included an additional basal lineage contribution to Qingdao, provided a significantly better fit than the two-way admixture model (Model 1) (p < 2.2 × 10^−16^, likelihood ratio test) (Fig. S5, Table S5). Although the estimated basal contribution was small (0.6%), its presence is further supported by introgression analyses (see below), suggesting that Qingdao retains traces of ancestry from an unsampled lineage.

### Dual origin of introgressed segments in the Qingdao population

To further investigate the origins of non-northern ancestry in the Qingdao population, we aimed to identify the putative introgressed segments from two distinct sources: one originating from southern populations and another from an unknown basal lineage. To identify these segments, we applied Skov HMM^47^, a model-free method for detecting the introgressed segments, for unphased genomes of the Qingdao population using two reference sets: (A) Korea and Okinawa and (B) an expanded set including Nanjing and Hangzhou (see Methods).

Qingdao genomes contain a substantial amount of introgressed segments detected using Reference A (66.9–74.3 Mbps per individual, 10.2–11.4% of the genome), with roughly 85% of private variants shared with Nanjing or Hangzhou and 15% not shared with any known populations (Fig. S7). Among these segments, those that did not overlap with those detected using Reference B were classified as “southern segments” (32.6–37.7 Mbps per individual), while overlapping segments were categorized as “ghost segments” (32.5–36.7 Mbps) (Table S6). These segments exhibit distinct genomic patterns characterized by PCA and *f_4_*-statistics: southern segments are enriched for genetic affinity with Hangzhou/Nanjing (Extended Data Fig. 1a–c), while ghost segments seem to be genetically unrelated to the southern populations (Extended Data Fig. 1c). QpGraph modeling shows that Qingdao receives 44% ancestry from a lineage predating the divergence of Korea-China populations in ghost segments, whereas Qingdao receives 27% gene flow from Nanjing in southern segments (Extended Data Fig. 1d– g).

Segment length distributions further support distinct admixture histories. Southern segments average 4,747 bp (4,569–4,883 bp; min–max of average length per individual) (Extended Data Fig. 2a), approximately half the length of ghost segments (8,120 bp; 7,808– 8,316 bp) (Extended Data Fig. 2b), suggesting older introgression. While mutation rates, estimated using nucleotide diversity from the Nanjing population, are slightly higher (5–10%) in introgressed segments due to stronger detection power in high-mutation regions (Extended Data Fig. 2c), both segment types exhibit 50-fold higher private mutation densities (4.3–4.4 mutations/kbp) than non-introgressed segments (0.08 mutations/kbp), supporting the authenticity of introgressed segments (Extended Data Fig. 2d–f).

The presence of southern and ghost ancestry in Qingdao is weakly but positively correlated with gene density (Spearman’s rank correlation coefficient = 0.06 and 0.05; p = 3.0 × 10^−3^ and 2.0 × 10^−2^, respectively) (Extended Data Fig. 3a–c), suggesting an absence of strong negative selection against the introgressed segments. However, both ancestry types are notably depleted in X chromosome-like scaffold 8, where segment lengths are also approximately two-fold shorter compared to the rest of the genome (Extended Data Fig. 3d,e).

Additionally, we observed a significantly lower ratio of heterozygosity between scaffold 8 and the whole genome among northern populations (Korea and Qingdao) compared to the others (Extended Data Fig. 4a). The median ratio of heterozygosity between scaffold 8 and the whole genome is 0.50 in Korea and 0.56 in Qingdao, in stark contrast to 0.98 in Hangzhou, 0.84 in Nanjing, and 0.91 in Okinawa. Under neutral conditions with an uneven sex ratio, the expected X-to-autosome heterozygosity ratio decreases from 1.1 to 0.6 as the male proportion in a population increases from 0 to 1. However, the observed values in Korea (0.50) and Qingdao (0.56) fall below this theoretical lower limit, implying that natural selection may have acted on scaffold 8 in northern populations (Extended Data Fig. 4b).

Interestingly, the ratio of introgressed proportion (southern + ghost) in scaffold 8 relative to the whole genome per individual is highly lower in male Qingdao P. longiforceps (0.02– 0.05) than females (0.48–0.63) (Table S6). Over 94% of introgressed variants found in the Qingdao population are shared by multiple individuals within the Qingdao or southern populations, indicating that this imbalance is not merely an artifact of lower male coverage on scaffold 8 (Fig. S8).

### Genomic signatures of positive selection during northward expansion

To identify genomic signatures of positive selection shared by the northern populations, we conducted genome-wide scans using two statistics: *f_4_* (Korea, Hangzhou; Qingdao, Nanjing) to assess intergroup genetic differentiation, and the nucleotide diversity ratio (π_Southern_/π_Northern_) to capture contrast in within-group diversity. We calculated them using a sliding window framework (50 kbp windows with 5 kbp steps) and after masking introgressed non-northern genomic segments in Qingdao. Of the 123,908 windows analyzed, 282 (0.2% of the genome) fell within the top 1% for both *f_4_* and π ratio (π_Southern_/π_Northern_), marking them as candidate regions under selection in the northern group (Fig. 5a, Table S7).

**Fig. 5.**
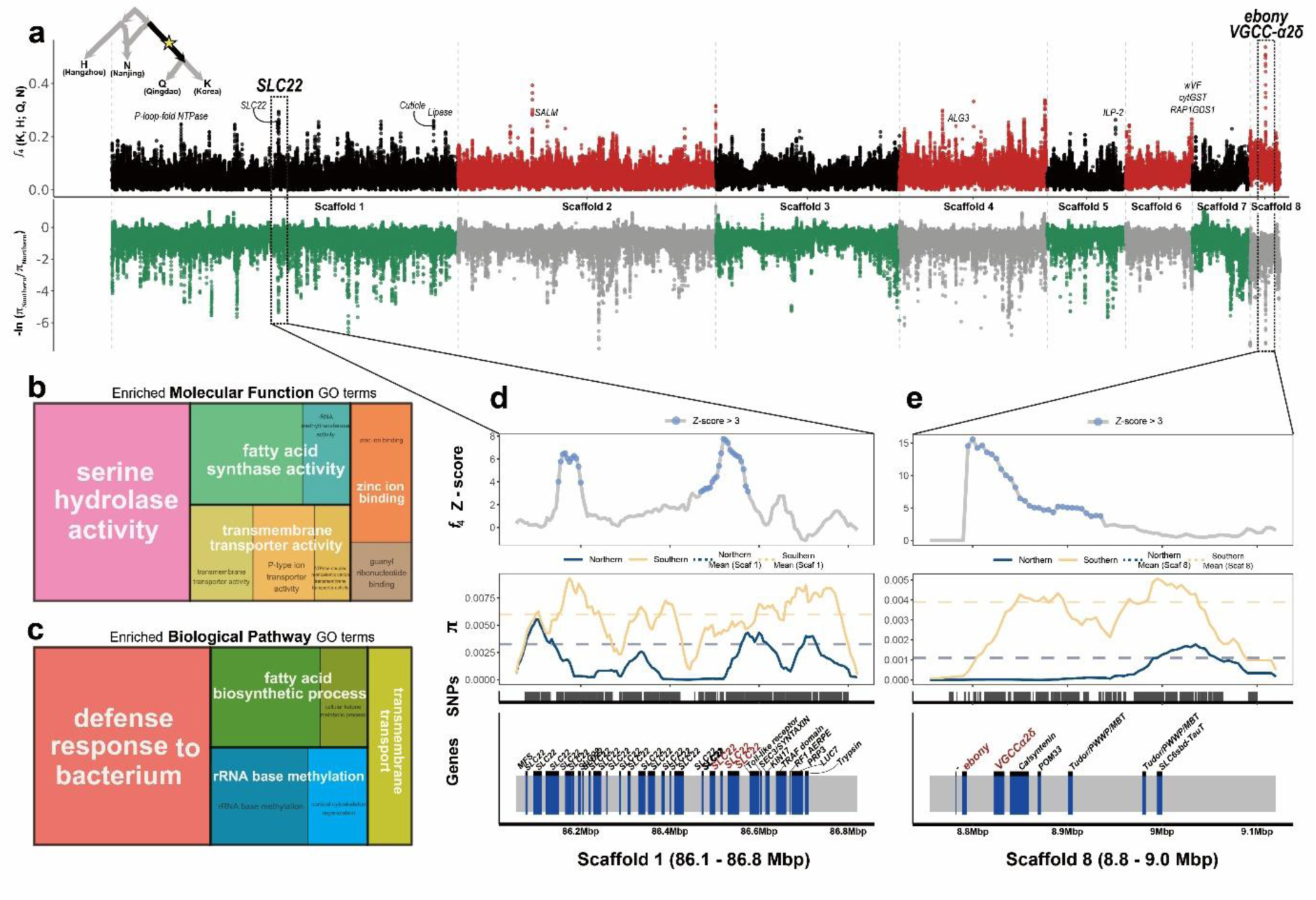
Genomic regions under positive selection in the northern group. **a**, Manhattan plots of *f₄*(Korea, Nanjing; Qingdao, Hangzhou) and the negative log-transformed nucleotide diversity ratio -ln (π_Southern_/π_Northern_) calculated in 50 kbp windows with a 5 kbp step size along scaffolds 1–8. Scaffolds are distinguished by alternating colors. Genes located within the top 1% of both *f₄* and π-ratio values were identified, and those within the 10 peaks with the highest *f₄* values are labeled. The top two genes showing the strongest signals are highlighted with dashed boxes. **b–c**, Treemap plots showing clustered and reduced Gene Ontology (GO) terms significantly enriched in candidate regions, defined as windows within the top 1% for both *f₄*and π-ratio, for **(b)** molecular function and **(c)** biological process. Enrichment was assessed using Fisher’s exact test with the “*weight01*” algorithm (p < 5.0 × 10^-2^). Size of the cluster indicates the sum of –log10 (*p*-value) within the cluster. **d–e**, Selective sweep patterns and the distribution of SNPs and annotated genes in the two regions with the strongest signals: **(e)** 8.8 – 9.0 Mbp on scaffold 8 and **(d)** 86.1 – 86.8 Mbp on scaffold 1. Z-scores for *f₄* were calculated using 1,000 randomly sampled windows, each at least 100 kbp apart. Horizontal dashed lines represent the scaffold-wide mean of π for each group. SNPs and gene annotations within the sweep regions are shown at the bottom.

Among these candidate regions, the strongest selection signal was detected on scaffold 8 at 8.8–9.0 Mbp (Fig. 5d) containing *ebony*, featuring the highest *f_4_* (*f_4_* top 0.001%, *π*-ratio top 0.05%), where the introgression is highly depleted in Qingdao individuals (Fig. S9). Notably, *ebony* is a key gene involved in cuticle formation, contributing to thermoregulation and desiccation resistance^48–50^. Its expression has been shown to vary along latitudinal clines in *Drosophila*, suggesting a role in climatic adaptation^51^. Alongside this, the region also contained *VGCC-α2δ* (voltage-gated calcium channel α2δ subunit), a temperature-sensitive neuronal signaling gene implicated in synapse formation and heat nociception^52–54^. Notably, thirteen fixed variants between northern and southern groups were located in the upstream region of *ebony*, between 1.9 kbp and 3.9 kbp from the gene. Additionally, five nonsynonymous variants in *VGCC-α2δ* exons showed pronounced allele frequency differences (northern = 0; southern = 0.92-0.94), suggesting potential functional divergence during northward expansion.

Another strong signal was identified on scaffold 1 at 86.6–86.8 Mbp (Fig. 5e), ranking second in *f_4_* signal strength (*f_4_* value top 0.03%, *π*-ratio top 0.4%). This peak is situated in the region harboring 23 copies of the *SLC22* (solute carrier protein family 22) gene, known to play a key role in management of oxidative stress induced by various environmental changes including cold stress, starvation, and immune activation^55,56^. The third strongest signal, located on scaffold 6 at 37.1–37.2 Mbp, included *vWF* (von Willebrand factor), a gene responsive to low temperature and implicated in immune and nutritional stress responses in arthropods^57,58^. Furthermore, an *ILP2* (Insulin-like peptide 2)-related gene, encoding a cold-induced insulin-like peptide with strong growth-promoting effects, showed the fourth strongest signal on scaffold 5 at 38.3–38.4 Mbp^59^.

Across all genes located within these candidate regions, the most significantly enriched GO term for molecular function was serine hydrolase activity (GO:0017171; p=5.2×10⁻^14^), which contributes to a wide range of physiological processes in insects including digestion, molting, and innate immune responses that are likely important for adaptation to novel environments (Fig. 5b)^60^. The most significantly enriched biological pathway term was defense response to bacterium (GO:0042742; p=2.5×10^−10^), suggesting an adaptive response to new bacterial exposures during range expansion (Fig. 5c)^61^. Furthermore, fatty acid synthesis and transmembrane transporter activity were highly enriched GO terms in both the molecular function (GO:0004312; p=1.2×10^−5^, GO:0022857; p=2.5×10^−3^) and biological pathway categories (GO:0006633; p=9.5×10^−4^, GO:0055085; p=4.0×10^−3^) (Fig. 5b, c). Notably, fatty acid synthesis is known as a key metabolism contributing to cold tolerance, energy storage, and desiccation resistance in various organisms through regulating components of membrane phospholipids and formulating triacylglycerols and cuticular hydrocarbons^62,63^.

### Lineage-specific positive selection in the Korea population

We next focused on selection signals specific to the Korean population not shared with Qingdao. We applied Population Branch Excess (PBE), a statistic derived from Population Branch Statistics (PBS) designed to reduce false positives^64^. In parallel, we calculated the nucleotide diversity ratio π_Qingdao_/π_Korea_ to assess reductions in genetic variation within the Korean lineage. Of the 119,979 windows analyzed, 262 (0.2% of the genome) fell within the top 1% for both PBE and π ratio (π_Qingdao_/π_Korea_), marking them as candidate regions under selection specific to the Korean population (Fig. 6a, Table S8).

**Fig. 6.**
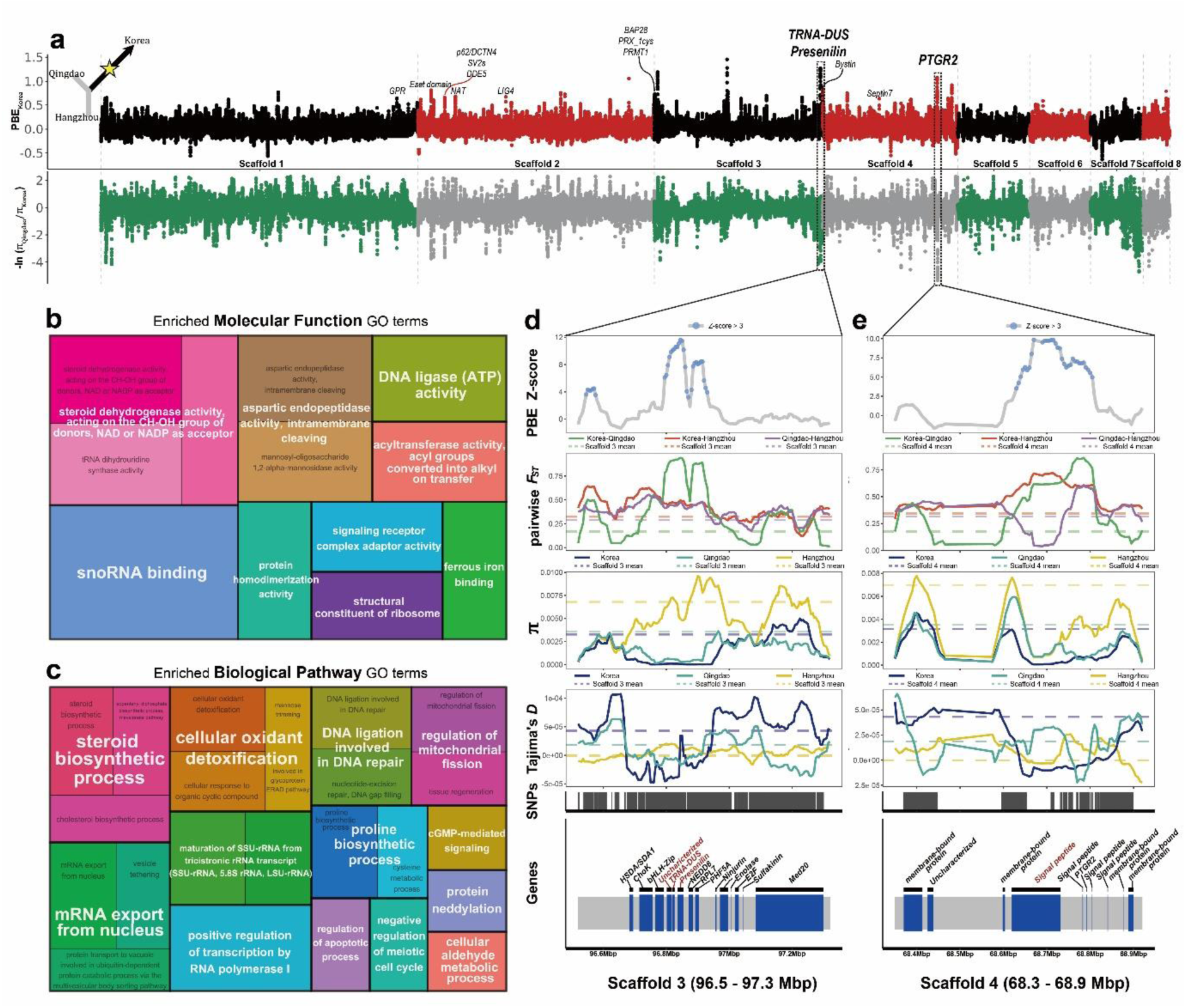
Genomic regions under Korea-specific positive selection. **a**, Manhattan plots of Population branch excess (PBE) of Korea and the negative log-transformed nucleotide diversity ratio -ln (π_Qingdao_/π_Korea_) calculated in 50 kbp windows with a 5 kbp step size along scaffolds 1–8. Scaffolds are distinguished by alternating colors. Genes located within the top 1% of both PBE and π-ratio values were identified, and those within the 10 peaks with the highest PBE values are labeled. The top two genes showing the strongest signals are highlighted with dashed boxes. **b–c**, Treemap plots showing clustered and reduced Gene Ontology (GO) terms significantly enriched in candidate regions, defined as windows in the top 1% for PBE and π-ratio, for (**b**) molecular function and (**c**) biological process. Enrichment was assessed using Fisher’s exact test with the “*weight01*” algorithm (p < 5.0 × 10^-2^). Size of the cluster indicates the sum of –log10 (*p*-value) within the cluster. **d–e**, Selective sweep patterns and the distribution of SNPs and annotated genes in the two genomic regions showing the strongest signals: (**d**) 96.5 – 97.3 Mbp on scaffold 3 and (**e**) 68.3–68.9 Mbp on scaffold 4. Horizontal dashed lines indicate the scaffold-wide mean of each corresponding statistic per group. Z-scores for PBE were calculated from 1,000 randomly sampled windows, each spaced at least 100 kbp apart. SNP positions and gene annotations within the sweep regions are displayed at the bottom of each panel.

The strongest selective sweep signal was detected on scaffold 3 at 96.8–96.9 Mbp (PBE top 0.004%; *π*-ratio top 1%) (Fig. 6d). This region included *Presenilin*, *Dus*, and one uncharacterized gene within the peak window, as well as *NEDD8* (Neural precursor cell expressed developmentally downregulated protein 8) and *RPL7* (Ribosomal Protein L7) immediately adjacent. *Presenilin*, enriched in the larval central nervous system, potentially promotes thermosensory neuron development via Notch-mediated neuronal differentiation^65,66^. *NEDD8* is known to protect against oxidative stress, likely associated with mitigating cold- and desiccation-induced damage^67^. In addition, *RPL7* has been shown to be upregulated during cold acclimation and freeze response in other taxa, suggesting a role as a physiological sensor of low temperature^68^.

Another highly significant selective sweep was detected on scaffold 4 at 68.6–68.8 Mbp, ranking second among candidate windows based on PBE values (PBE top 0.02%; π-ratio top 0.3%) (Fig. 6e). This region contains *PTGR2* (Prostaglandin reductase 2), a key enzyme involved in the catabolic pathway of prostaglandins, along with four unspecified signal peptides and two membrane-bound proteins. Prostaglandins are bioactive lipids that influence reproduction, immune responses, and melanization in insects, and are also implicated in thermal homeostasis in mammals^69,70^. Notably, experimental evidence in Lepidoptera has shown that suppressed expression of *PTGR* leads to prolonged immune activation and increased melanization, resulting in darker larval pigmentation^71^. A variant in the intronic region of *PTGR2* (position 68,791,127 on scaffold 4), absent in both Hangzhou and Qingdao but present at a high frequency of 0.92 in the Korean population, highlights potential expressional changes in this gene during local adaptation. Both of the top two genomic signals were further supported by significantly reduced Tajima’s *D* values in the Korean population, ranking in the bottom 0.01% and 0.05% respectively, consistent with expectations under recent selective sweeps.

The Gene Ontology enrichment analysis of these candidate regions of Korea specific positive selection revealed significant overrepresentation of sterol metabolism genes in both molecular function (GO:0033764; p=1.9×10^−2^) and biological process categories (GO:0006694; p=1.2×10^−2^, GO:0006695; p=2.5×10^−2^) (Fig. 6b, c). Given its roles in membrane fluidity regulation and molting hormone production, sterol metabolism is critical in insects, and their dependence on dietary sterols may drive selection on sterol-processing pathways in unfamiliar habitats^72,73^. Additionally, the enrichment of oxidative stress response pathways (GO:0051920; p=3.7×10^−2^, GO:0098869; p=1.9×10^−2^) and DNA repair mechanism (GO:0051103; p=1.8×10^−2^, GO:0006297; p=2.0×10^−2^, GO:0003910; p=1.8×10^−2^) possibly align with the need to cold/UV stress resilience mechanisms in shifted environment^74,75^.

## Discussion

We present population-scale genome data of *P. longiforceps* across major regions in East Asia, providing strong evidence for the distinct origins of Korea and Okinawa *P. longiforceps* populations. Previous studies proposed long-distance translocations from the southern, subtropical regions to Korea, possibly associated with human activities^28^. Based on our discovery of *P. longiforceps* in Qingdao, and the close genetic distance between Qingdao and Korea among all sampled populations, we suggest that Korean *P. longiforceps* may instead have arrived from a northern source. The Shandong peninsula is not only closer to Korea, but also hosts several major harbors (e.g. Qingdao) with active trade between Incheon port near Seoul. Qingdao, however, has additional genetic affinity to southern populations, and divergence from Korean populations is estimated to have occurred at least 900 generations ago, well before the swarming of *P. longiforceps* was observed in Korea. Considering the pattern of isolation-by-distance along the geographical gradient, the Korean populations likely originate from a yet undiscovered northern lineage, possibly in the northeastern regions of Shandong, even farther north than Qingdao. On the other hand, the Okinawa population is genetically intermediate between Nantou and Pingtung, and thus neither of the sampled Taiwanese populations serves as direct sources for Okinawan individuals. As an alternative, we can infer the contribution of another source population from underrepresented regions such as northern Taiwan, which would be geographically closer to both the Ryukyu islands and mainland China.

The population structure of continental *P. longiforceps* suggests multiple northward migrations over an extended time scale. We identified two distinct sources of introgression in Qingdao individuals: one (“southern”) aligns closely with Nanjing (Fig. S7B) and the other originates from an unidentified (“ghost”) lineage split slightly earlier than the first split among all sampled continental *P. longiforceps* populations, suggesting a metapopulation structure (Extended Data Fig. 1g). Similar mutation densities of southern and ghost segments indicate multifurcation rather than deep divergence (Extended Data Fig. 2d,e). Historical records hint at Jiangxi and Fujian^26,76^ as possible locations of the metapopulation, although ghost ancestry may have arrived via intermediates. The length distributions of both southern and ghost segments follow an exponential decay, with ghost segments being roughly twice as long as southern segments (Extended Data Fig. 2a,b). This pattern supports at least two independent pulse-like admixture events. Notably, we did not detect any introgressed segments longer than 207 kbp, providing a lower bound for admixture timing. This pattern suggests that the dispersal from southern to northern China was sporadic rather than continuous.

Population bottlenecks are a common feature of biological invasions, particularly when a small number of individuals establish a new population. Long-term demographic projections and severely reduced genetic diversity in northern populations suggest serial founder events during the northward expansion. Furthermore, the invasive *P. longiforceps* population in Korea exemplifies successful colonization despite an extreme recent bottleneck, with its effective population size shrinking to approximately 100 individuals. Despite our extensive sampling efforts, Korean specimens held an extreme level of genetic homogeneity compared to native populations in southern China. Thus, a single introduction of a small group seems the most parsimonious scenario for the origin of *P. longiforceps* in Korea. A similar pattern of genetic homogeneity and severe bottleneck in Okinawa suggests that the recent invasion success of *P. longiforceps* in both regions has not, thus far, been driven by repeated introduction events or admixture with native populations.

As *P. longiforceps* migrated toward higher latitudes, shifts in seasonal patterns of temperature likely played a major role in their adaptation. Mating swarms rely on synchronized emergence for successful reproduction, especially within the tight seasonal windows of northern populations^28^. The strongest signature of selection in northern populations was identified in the 8.8–9.0 Mbp region of scaffold 8, which encompasses the genes *VGCC-α2δ* and *ebony*, associated with thermal sensing and regulation, respectively^77–79^. Selection on these loci may have facilitated synchronization of reproductive cycles with seasonal changes and improved thermal resilience.

Scaffold 8 overall shows a significant reduction of genetic diversity in northern populations, and segments of introgression on scaffold 8 are reduced both in number and size. This implies that strong purifying selection has been purging non-northern alleles. Furthermore, the heterosomal properties of scaffold 8 suggests that this process may be sex-biased. The coverage ratio of scaffold 8 relative to the whole genome of males and females (median; male = 0.6, female = 1.1) suggests that *P. longiforceps* has an XO sex determination system like other bibionids, with scaffold 8 likely being the X chromosome (Table S1)^80,81^. Male *P. longiforceps* in Qingdao have especially low levels of introgressed proportions on scaffold 8, only a tenth of that in females. This indicates that ongoing purifying selection is occurring faster and stronger in Qingdao males, possibly due to the smaller effective population size, biased expression levels, or sex-specific function and selection of X-linked loci in hemizygous individuals.

The selection signals shared within the northern group and specific to Korea indicate recurrent positive selection in pigmentation, lipid metabolism, immune system, and stress response. Notably, these functional categories are well documented in *Drosophila* as being associated with climatic and environmental variation. In particular, pigmentation is one of the most well studied traits exhibiting clear latitudinal clines in *Drosophila*^51,82^. Comparable genetic patterns in *P. longiforceps* may similarly indicate adaptation along a geographic gradient. Latitudinal divergence in *Drosophila* has also been observed in lipid uptake allele expression, total lipid content, immune gene expression, and indel variation in oxidative stress-related genes^83–85^.

To further refine these findings, additional efforts in sample collection, phenotypic data acquisition, and ecological surveys are required. Expanded sampling from northern China may help pinpoint the direct source of the Korean population and clarify whether adaptive changes occurred before or after its arrival in Korea. Comparative studies including insular populations will be particularly valuable for understanding how this species adapt across different temporal scales. Large-scale phenotypic surveys, particularly of lipid content and pigmentation, are required to validate the functional impact of selection signals and delineate the direction of selection acting on northern populations. Finally, investigating how the dispersal and outbreak dynamics of this species affect native ecosystems represents an important direction for future ecological research. As the number of species introduced into new environments continues to rise, the need for comprehensive studies on invasive species grows accordingly. Continued research on *P. longiforceps* will provide valuable insights for understanding the evolutionary dynamics of biological invasions.

## Methods

### Sample provenance

A total of 156 individuals of *P. longiforceps* were sampled from 32 sites in seven major geographical regions across its range in East Asia (Table S1). Samples from Korea, Japan, and China were collected between 2021 and 2024, and preserved in 99% ethanol at −20 °C or packed with dry ice until further procedures. Samples from Taiwan were acquired from the National Museum of Natural Science, Taichung, Taiwan.

### Laboratory procedures

For samples from Korea, Japan, and Taiwan (n=118), genomic DNA was extracted using the OmniPrep for Tissue kit (G-Biosciences, USA). DNA extracts from Taiwan samples were processed using the REPLI-g Single Cell Kit (Qiagen, USA) for whole genome amplification prior to library construction. Illumina sequencing libraries were prepared according to the QIAseq FX DNA Library Kit (Qiagen, USA) protocol. In brief, 100 ng of genomic DNA were fragmented using the FX Enzyme Mix for 10 minutes at 32°C, followed by ligation of Illumina adapters. The resulting product was purified with AMPure beads, performing two rounds of bead-based selection: 80 µL of beads for the first round and 50 µL for the second round. The purified product was then PCR-amplified and size-selected for fragments between 450 and 600 bp using the PippinHT system (Sage Science, USA). Finally, 150 bp paired-end sequencing was performed on HiSeq X Ten or NovaSeq 6000 platforms (Illumina, USA) by Macrogen (Seoul, Korea). Genomic DNA extraction, library preparation, and sequencing for samples from China (n=38) were conducted by Novogene (Tianjin, China). Illumina libraries were prepared using the NEBNext Ultra II DNA Library Prep Kit (New England Biolabs, USA) and 150 bp paired-end sequencing was performed on a NovaSeq X Plus platform (Illumina, USA).

### Genome data processing

The sequence reads were aligned to the *P. longiforceps* reference genome assembled from a Korean sample^34^ employing bwa mem v0.7.17^86^ with default parameters. Following alignment, the resulting BAM files were sorted and indexed using Samtools v1.9^87^. Potential PCR duplicates were identified utilizing the MarkDuplicates tool of Picard v2.20.2 (available at http://broadinstitute.github.io/picard). We filtered out reads with mapping quality less than 30 using Samtools v1.9^87^.

For the variant calling from the BAM files, a GVCF (Genomic Variant Call Format) file was generated using the HaplotypeCaller module in GATK with the “-ERC GVCF” option^88^. Individual GVCF files were merged using the CombineGVCFs module in GATK, excluding 6 samples with low endogenous DNA (<40%) or high bacterial contamination (>25%), estimated using Kraken v2.1.3 and minikraken2_v1_8GB^89^. SNPs were then called and selected from the combined GVCF file using the GenotypeGVCFs and SelectVariants modules, respectively. To minimize false-positive variants, low-quality variants were filtered out using the VariantFiltration module of GATK when a variant meets at least one of the following criteria: (1) SNPs with mean DP (for all individuals) lower than 1/5 or higher than 5 times of the genome-wide average across all SNP sites, (2) QD (Quality by Depth) < 2, (3) phred-scaled variant quality score (QUAL) < 30, (4) SOR (Strand Odds Ratio) > 3, (5) FS (Fisher Strand) > 60, (6) MQ (Mapping Quality) < 40, (7) MQRankSum (Mapping Quality Rank Sum test) < −12.5, (8) ReadPosRankSum (Read Pos Rank Sum test) < −8, and (9) ExcessHet > 10. Subsequently, variants other than biallelic SNPs were excluded, resulting in 44,877,370 high-quality biallelic SNPs. Each variant was genotyped for all samples used for variant calling (n=150) based on its genotype likelihood. The most likely genotype for each individual and variant was taken if its normalized likelihood was 0.9 or more; otherwise, the genotype was treated as missing data.

Considering the heterogeneity of sequencing depths of samples and large genetic distances between Taiwan-Okinawa populations and reference genome from Korean population, we call pseudo-haploid genotypes for all samples used for variant calling (n=150) by following procedures: one high-quality base (Phred-scaled base quality score ≧ 30) was randomly sampled using pileupCaller v1.5.2 (https://github.com/stschiff/sequenceTools) for each of the 44.9 million SNPs discovered from the genome data processing.

To exclude variants from low-quality genomic regions, we removed SNPs within genomic windows of the *P. longiforceps* reference genome that met one or more of the following criteria: (1) 35 bp windows with low mappability (c < 3), following Heng Li’s 35mer filter with r=0.5 (https://lh3lh3.users.sourceforge.net/snpable.shtml), (2) 1 kbp windows that had extreme GC content (top 2.5% or bottom 2.5%); (3) 1 kbp windows with mean depth < 1/5x or > 5x (x: overall mean sequencing depth across all 1kbp windows) in at least one of Korea, Hangzhou, and Okinawa; (4) 1 kbp windows with high repeat content (> 90%); and (5) 1 kbp windows that included one or more undefined bases in the reference genome (N/n). Furthermore, we disregarded SNPs with ≧5 missing individuals or minor allele counts ≦2 in the pseudo-haploid genotypes set. The resulting set of 16,989,044 SNPs for both diploid and pseudo-haploid genotypes were utilized for subsequent analyses.

To assess whether variations in sequencing depth and differences in genetic distance to the reference genome significantly impact analyses based on diploid genotypes, we compared principal component analysis (PCA) and *f*-statistics using both diploid and pseudo-haploid genotype data. We found no clear inconsistencies between PCA and f-statistics when using pseudo-haploid or diploid genotypes (Figs. 2a and S10; Tables S3, S9). Since diploid genotype calls retain more information, we used diploid genotypes in subsequent analyses to achieve higher resolution.

### Principal component analysis

For PCA, we calculated eigenvectors with 150 *P. longiforceps* diploid and pseudo-haploid genotypes using smartpca v.18140^90^ with the option ‘lsqproject: YES’.

### F-statistics analysis and qpGraph

We employed the “qp3pop”, “fst”, and “qpdstat” functions from the ADMIXTOOLS2 package^91^ to compute outgroup *f*_3_ and *f*_4_-statistics. To estimate genetic differentiation, we calculated Hudson’s *F* ^92^ between all population pairs, excluding Taiwan due to low sample size. We computed *f*_4_ statistics with the “f4mode = TRUE” option. We used the Okinawa population for all f-statistics as an outgroup for Chinese and Korean *P. longiforceps* populations, while the Korean population served as an outgroup for the Taiwan and Okinawa populations. We used *f*_4_ statistics in the form of *f*_4_(Outgroup, X; target1, target2) to test symmetry between targets or search for additional sources, and *f*_4_(Hangzhou/Nanjing, Qingdao/Korea; island 1, island 2) to assess asymmetry between continental and island populations.

We employed the “qpgraph” function from the ADMIXTOOLS2 package^40,91^ to identify the most plausible admixture graph topologies of Korea-China populations employing 5 populations: Okinawa, Hangzhou, Nanjing, Qingdao, Korea. We used 16,927,249 SNPs for qpGraph where all 5 populations have genotypes of at least one chromosome.

### Two-taxon gene flow test

We utilized two-taxon A_m_ statistics to test whether a gene flow event happened between island and continental populations^93^. Following Mackintosh and Setter 2024^93^, A_m_ statistics between individual A and B in specified block size L bp is defined as following formula:

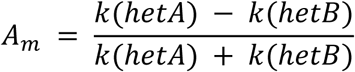

where k(hetA) and k(hetB) are the counts of heterozygous calls of the individual A and B, respectively observed less than L bp from the site where both A and B are heterozygotes. We counted k(hetA) and k(hetB) only when both individuals have non-missing calls on the site. We used a single genome with the highest coverage for each continental (Korea, Qingdao, Nanjing, and Hangzhou) and island populations (Okinawa, Nantou). We calculated A_m_ statistics for every possible pair of continental and island populations for a range of block sizes from 2 to 128 bp. We restricted analysis in small block sizes to avoid confounding effects due to recombination disrupting linkage. We estimated a standard error using 5 Mb block-jackknifing.

### Heterozygosity, Run of homozygosity (ROH), and identity by descents (IBD)

We estimated genome-wide heterozygosity for each sample with coverage greater than 8× (n=137) by counting the total number of heterozygous biallelic SNPs from the final VCF sets using VCFtools v0.1.16 with the “--het” option^94^. This sum was divided by the total number of callable nucleotide positions in the individual genome, calculated using qualimap v2.1.1^95^. For a single Nantou genome (TW316), we estimated a standard error using 10 Mb block-jackknifing.

To identify long ROH blocks, we applied a hidden Markov model classification program in BCFtools v1.19^96^, assuming a conservative recombination rate of 1 cM/Mb. This rate is two or three-times lower than *Drosophila* recombination rates, which range from 2.06 to 3.44 cM/Mb^44^. We grouped (Korea, Qingdao), (Nanjing, Hangzhou), and Okinawa populations separately to calculate population-specific allele frequencies, as recommended by BCFtools. ROH blocks were called using the following command with default parameters: “bcftools roh -e --S $GROUP.list.txt -O r $VCF -o $GROUP.roh”. From each output file, we selected ROHs of at least 500 kbp in length with quality scores of ≥ 70.

IBD segments were detected from PLINK files of all *P. longiforceps* genomes (n=137, coverage > 8×) using IBIS v1.20.3^97^ with parameters “-mt 300” and “-er 0.004”. We assumed a conservative recombination rate of 1 cM/Mb and used a minimum shared segment length threshold of 7 cM to minimize false-positive detections.

### Population size history

We estimated changes in the effective population size (N_e_) of *P. longiforceps* over the past 100 generations using GONE^43^ (Santiago et al. 2020). Unlike coalescent-based methods (e.g., PSMC, MSMC), GONE leverages linkage disequilibrium patterns in phased or unphased genomes and performs well in estimating population size over recent hundreds of generations. We ran GONE on unphased diploid genotypes from Korea, Qingdao, Nanjing, Hangzhou, and Okinawa populations, selecting SNPs from scaffolds 1-7 to ensure sufficient SNP counts while excluding shorter scaffolds and the sex chromosome (scaffold 8). For each population, we performed independent runs using randomly selected SNP subsets (50,000 polymorphic SNPs per scaffold) and assumed a conservative recombination rate of 1 cM/Mb to prevent overestimation of recent bottleneck sizes.

For long-term Ne trends, we applied MSMC2 v2.1.1^45^ to individuals with the highest genome coverage from Korea, Qingdao, Nanjing, Hangzhou, and Okinawa populations. A mappability filter based on Heng Li’s SNPable filter was applied, retaining uniquely mappable regions (c=3) (http://lh3lh3.users.sourceforge.net/snpable.shtml). Additional filtering removed sites with coverage below half or above twice the mean scaffold coverage using bamCaller.py from msmc-tools. MSMC2 was run using scaffolds 1-7, excluding scaffold 8 and shorter scaffolds, with a within-individual coalescence model (-I 0-1,2-3), 10 bootstraps, and the time segment pattern “-p 1*2+25*1+1*2+1*3”. We split the first time segment for Okinawa genomes using the parameter “-p 27*1+1*2+1*3” as we observed a false peak and sudden decline with default parameters, as noted by Hilgers et al. 2025^98^. Population size curves were scaled using a mutation rate of 5.0 × 10⁻⁹ per generation per base pair.

For X chromosome pseudodiploid analysis, we used one high-coverage male per population on scaffold 8. SNP calling followed the same filtering criteria as autosomal MSMC2, with additional parameters for haploid male X chromosomes (“bcftools call -m -f GQ,GP -- ploidy 1”). MSMC2 was run on cross-population pairs to estimate the coalescence rate of pseudodiploid between a pair of populations. Finally, the combineCrossCoal.py script from the msmc-tools was used to plot the relative cross-coalescence rate (rCCR) using the following calculations:

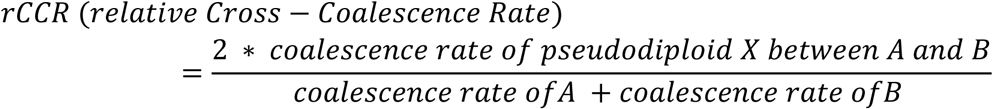

Mutation rate scaling was omitted due to population-specific autosome/X diversity ratios. Population divergence was estimated at the left end of the most recent time interval where rCCR ≥ 0.5^99^.

### Explicit demography modelling

We used Momi2 (v.2.1.19)^46^ to model the demographic history of Korea-China populations. Two highest-coverage individuals per population were selected, and SNP sites were filtered using criteria from genome data processing: (1) 35 bp windows with low mappability (c<3), following Heng Li’s 35mer filter with r=0.5 (https://lh3lh3.users.sourceforge.net/snpable.shtml), (2) 1 kbp windows that had extreme GC content (top 2.5% or bottom 2.5%); (3) 1 kbp windows with mean depth < 1/5x or > 5x in one of Korea, Hangzhou, and Okinawa; (4) 1 kbp windows with high repeat content (> 90%); and (5) 1 kbp windows that included one or more undefined bases in the reference genome (N/n). Ancestral alleles were polarized using consensus genotypes from the Okinawa population. The unfolded site frequency spectrum (SFS) was split into 100 blocks for jackknifing and bootstrapping. The backbone tree topology was set as (((Korea, Qingdao), Nanjing), Hangzhou), with additional gene flow from Hangzhou into Nanjing. We tested alternative two-pulse migration models into Qingdao from (1) Nanjing, (2) Hangzhou, and (3) a basal lineage branching off all Korea-China populations. Ten independent runs with randomized starting parameters were performed, selecting the model with the highest log-likelihood. The best model underwent 100 bootstraps for confidence intervals.

### Detection of introgressed segments

To detect introgressed segments in Qingdao *P. longiforceps*, we applied the Skov Hidden Markov Model (HMM)^47^. Two reference panels were used to filter private variants:

- Reference A: Korean and Okinawa *P. longiforceps*, removing shared northern-derived variants and private ancestral variants in Qingdao.
- Reference B: Reference A plus Nanjing and Hangzhou, filtering shared derived variants within southern populations.

HMM was run with default parameters and a 1,000 bp window size. Callable regions followed Momi2 filtering criteria, and introgressed segments were detected using a posterior probability cutoff of 0.8^47^. Introgressed segments detected using Reference A that overlapped with those detected using Reference B were categorized as “ghost”’ segments, whereas non-overlapping segments detected using Reference A were classified as “southern” segments. To infer introgression sources, we calculated *f*_4_-statistics and assessed affinity with Nanjing/Hangzhou using ADMIXTOOLS2^91^. We further applied qpGraph^40^ to fit Qingdao under different introgression scenarios (from Nanjing, Hangzhou, or a basal Korea-China lineage). The frequency of introgressed segments was estimated per 250 kbp window using BEDTools v2.31^100^, and we tested correlations between segment frequency and gene density using Spearman’s rank correlation in R.

### Selection scan

We calculated multiple genome-wide statistics to robustly identify genomic regions under positive selection. We applied a sliding window approach with window size 50 kbp and step size 5 kbp), removing windows containing fewer than 100 SNPs. To establish genome-wide background distributions for each statistic, we randomly sampled 10,000 windows separated by at least 100 kbp, and calculated mean, median, standard error, and Z-scores where applicable. To detect signatures of selection related to northward expansion, we defined Hangzhou and Nanjing as the southern group and Qingdao and Korea as the northern group, then measured genetic differentiation between groups as well as genetic diversity within groups. We calculated *f_4_*-statistics of the form *f_4_*(Korea, Nanjing; Qingdao, Hangzhou) to quantify internal branch lengths between southern and northern groups, identifying regions with excess differentiation. We also estimated nucleotide diversity (π) for each group and computed π-ratio (π_Southern_/π_Northern_). Candidate selective sweep regions were selected as genomic windows ranking in the top 1% for both *f_4_* and π-ratio, and we prioritized these windows based on their *f_4_* values.

To identify positive selection signals specifically in the Korean population, we employed the population branch excess (PBE) statistic, a derivative of population branch statistics (PBS)^101^ designed to reduce false positives^64^. PBE measures the relative elongation of a focal population’s branch compared to total branch length across three populations, thus effectively identifying rapidly evolving regions while accounting for confounding scenarios such as balancing selection, genetic incompatibility, or parallel selective sweeps^102^. To calculate PBE, we first computed window-specific PBS values for Korea and branch lengths between Qingdao and Hangzhou (T_Qingdao-Hangzhou_) using Hudson’s estimator of *F* ^92^. We then calculated genome-wide medians of these values, deriving window-level PBE using the following equations:

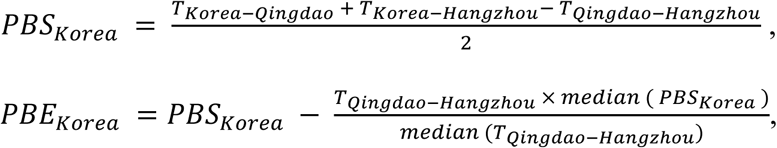

where *T* represents the relative divergence time between each pair of populations, estimated using the formula

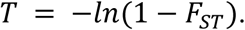

We estimated nucleotide diversity (π) for each group and calculated the π-ratio (π_Hangzhou_/π_Korea_). In addition, we computed Tajima’s D for each window to assess deviations from neutrality and infer the direction of selection^103^. We defined candidate selective sweep regions specific to Korea as windows simultaneously ranking in the top 1% for PBE and π-ratio and subsequently prioritized these windows according to their PBE values.

To distinguish independent signals and reduce redundancy, we conducted a window collapsing procedure based on local peaks. We identified the range of each local peak by detecting changes in the slope of each statistic, and then merged adjacent windows around each peak into single collapsed regions. To minimize false positives of *f_4_* and PBE resulting from introgression, we masked introgressed genomic segments in the Qingdao population using Skov’s HMM before performing subsequent analyses. We identified genes within candidate selection regions and matched them with functional annotations. If collapsed regions contained one or more genes, we retained these genes and retrieved their annotations, explicitly noting any gene located within peak windows. If no genes were identified within a collapsed region, we selected the nearest gene. We compiled final gene lists based on annotations associated with the lowest E-value. Gene prediction was performed using BRAKER3 v.3.0.8^104^, incorporating the OrthoDB v.12 Arthropoda protein dataset^105^ and RNA-seq data from *P. longiforceps*^34^. We further annotated predicted protein functions using InterProScan-5.65-97.0^106^ and assessed variant effects with ANNOVAR^107^ and SnpEff^108^ for comprehensive variant interpretation.

### Gene ontology enrichment analysis

Beyond examining individual top-signaling genes of positive selection, we conducted Gene Ontology (GO) enrichment analyses to identify molecular functions and biological pathways significantly enriched among candidate genes associated with adaptation. We retrieved functional annotations and corresponding GO terms for genes located within the candidate regions selected in the previous step as under positive selection in the northern group and specifically within the Korean population, respectively.

We constructed a custom annotation database (orgDB) for *P. longiforceps* using *AnnotationForge* v.1.48.0^109^. For each candidate gene set (Northern and Korea-specific), we performed GO term enrichment analysis in Molecular Function (MF) and Biological Process (BP) categories using Fisher’s exact test implemented in *topGO* v.2.58.0 with the ‘*weight01*’ algorithm^110^. In downstream analyses, we used *p*-values generated by the *weight01* algorithm without applying additional multiple testing correction, in accordance with the original recommendations of the library authors. We computed semantic similarity among significantly enriched GO terms (p < 5.0 × 10^−2^), clustered these terms to reduce redundancy using the *GOSemSim* v.2.32.0 package^111^, and visualized representative GO terms using the *rrvgo* v.1.18.0 package^112^.

## Data availability

All data needed to evaluate the conclusions in the paper are present in the paper and/or the Supplementary Materials. All newly generated sequencing data reported in this study, including raw reads (FASTQ) and aligned reads (BAM), are available from the European Nucleotide Archive under the accession number PRJEB87434.

## Code availability

This study makes use of publicly available software, referenced throughout the main text and supplementary material.

## Acknowledgements

We thank S. Dowell and M. Dong for their help in collecting samples from Okinawa, and W. J. Bang, S. Han, W. Kim, W. Park, J. Chung, and K. Kim for their assistance in collecting Korean specimens. We also thank J.-F. Tsai and B.-C. Lai for their help in acquiring samples from Taiwan. This study was supported by grants from the National Institute of Biological Resources (NIBR) funded by the Ministry of Environment (MOE) of the Republic of Korea (No. NIBR202410201, NIBR202505104, NIBR202510201), National Research Foundation of Korea (NRF) grants funded by the Ministry of Science and ICT (No. RS-2021-NR060088, RS-2021-NR061223), the Global-LAMP Program of the NRF grant funded by the Ministry of Education (No. RS-2023-00301976 to C.J.), and the Hyper-Convergence Research Program (No. 0525-20240128) and Creative-Pioneering Researchers Program through Seoul National University (SNU) (to C.J. and S.S.).

## Author Contributions

S.S. and C.J. conceived and supervised the study. Donghee Kim, J.C., and Dongyoung Kim curated and analyzed data. J.C., Dongyoung Kim, S.K., C.L.T., M.J.B, S.J.P, A.B., and S.S. provided samples for analysis. Donghee Kim, J.C., Dongyoung Kim, S.S. and C.J. wrote the manuscript with the input from all the other co-authors. Donghee Kim, J.C., Dongyoung Kim, M.C.L., S.S. and C.J. revised the manuscript.

---

**Extended Data Fig. 1.**
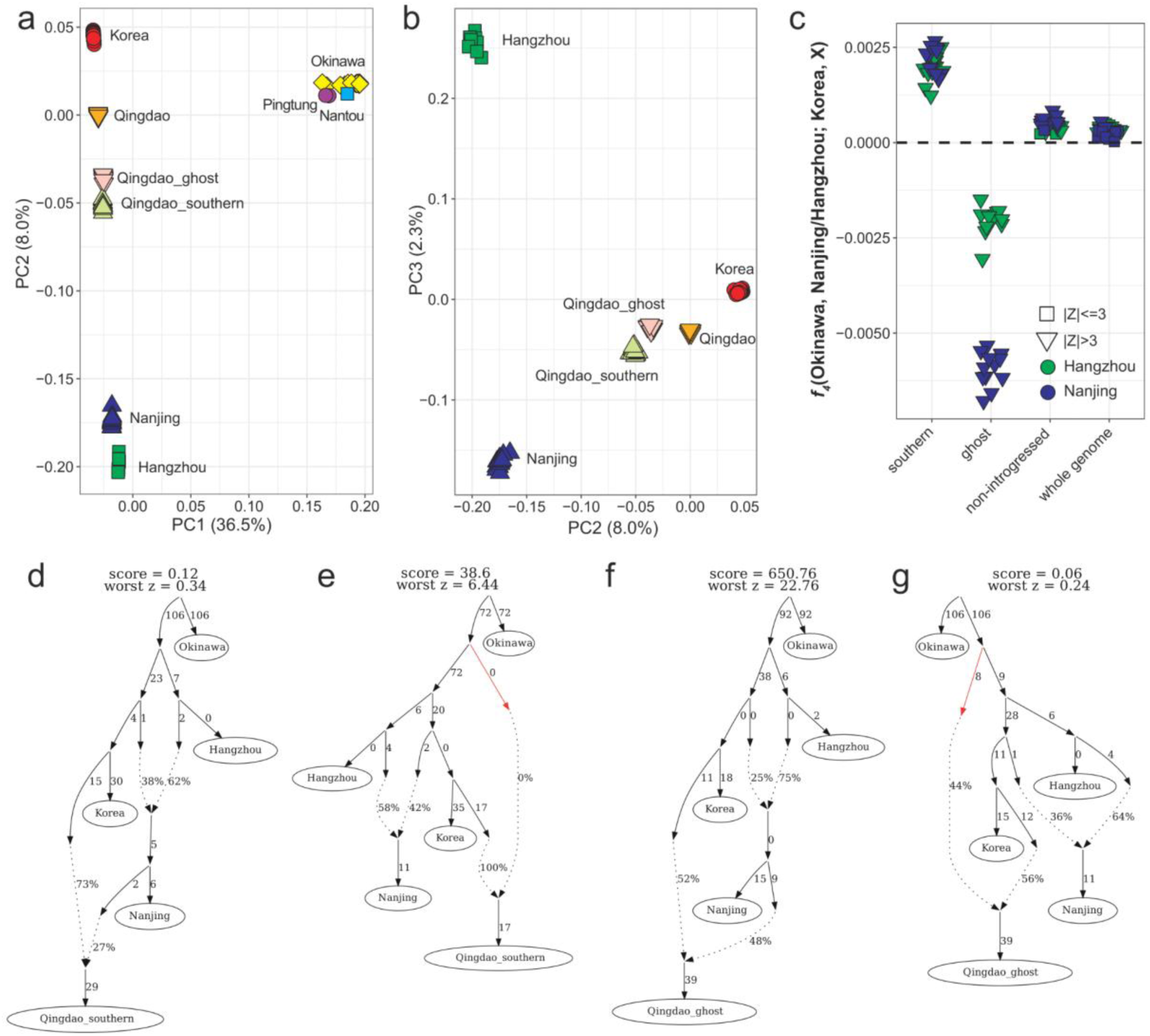
Two distinct origins of introgressed segments in the Qingdao *P. longiforceps* genome. **a**–**b**, Principal component (PC) plots of PC1 vs. PC2 (**a**) and PC2 vs. PC3 (**b**) based on the genotypes of 150 East Asian *P. longiforceps*. Qingdao individuals’ genotypes from non-introgressed, ghost, and southern segments are projected onto these PCs. Each individual is represented by a color-filled symbol corresponding to its group. The proportion of variance explained by each principal component is indicated on the axis labels. **c**, *f*_4_-statistics of the form *f*_4_(Okinawa, Nanjing/Hangzhou; Korea, X), computed for each Qingdao individual using genotypes from southern, ghost, non-introgressed segments, and the whole genome. Each point represents an estimate, with colors indicating the southern population used in *pop2*: green for Nanjing and blue for Hangzhou. The shape of each point reflects the significance of the Z-score: downward triangles for |Z| > 3 and circles for |Z| ≤ 3. Standard errors were calculated using a 5 cM block jackknife. **d**–**e**, Population graph fits modeling a gene flow event from Nanjing into Qingdao (**d**) and from a basal population into Qingdao (**e**), using genotypes from southern segments in Qingdao individuals. **f**–**g**, Same as **d**– **e**, but using genotypes from ghost segments. **d**–**g**, edge lengths correspond to *F_ST_* × 1000. The graph’s log-likelihood score and the largest residual deviation between observed and expected *f*-statistics (“worst *z*”) are displayed above each graph.

**Extended Data Fig. 2.**
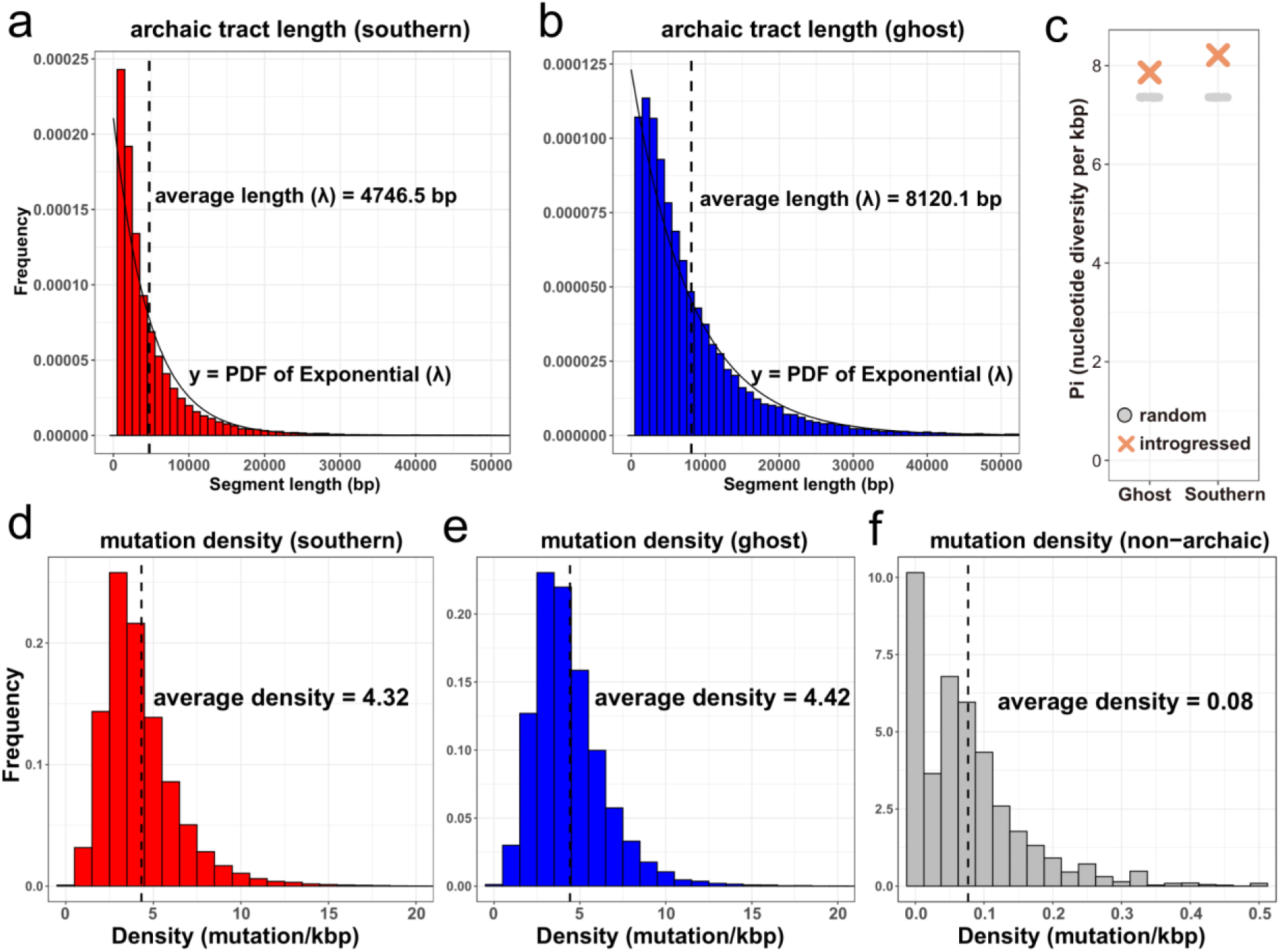
Length distribution and mutation density of introgressed fragments. **a**, Length distribution of southern segments. The average segment length is indicated next to the vertical dashed line. An exponential distribution fitted to the data, with the parameter set to the average length, is shown as a solid line. The x-axis ranges from 0 to 50,000 bp. **b**, Same as **a**, but for ghost segments. **c,** Nucleotide diversity (π) of Nanjing populations within introgressed (southern and ghost) segments compared to random regions matched for mean and total length. Random regions are sampled for 500 times each. π values for introgressed segments are shown as colored crosses, while those for random regions are marked as gray circles. **d**, Distribution of mutation density per 1-kbp window in southern segments. The average mutation density is annotated next to the vertical dashed line. **e**, Same as **c**, but for ghost segments. **f**, Same as **c**, but for non-introgressed segments.

**Extended Data Fig. 3.**
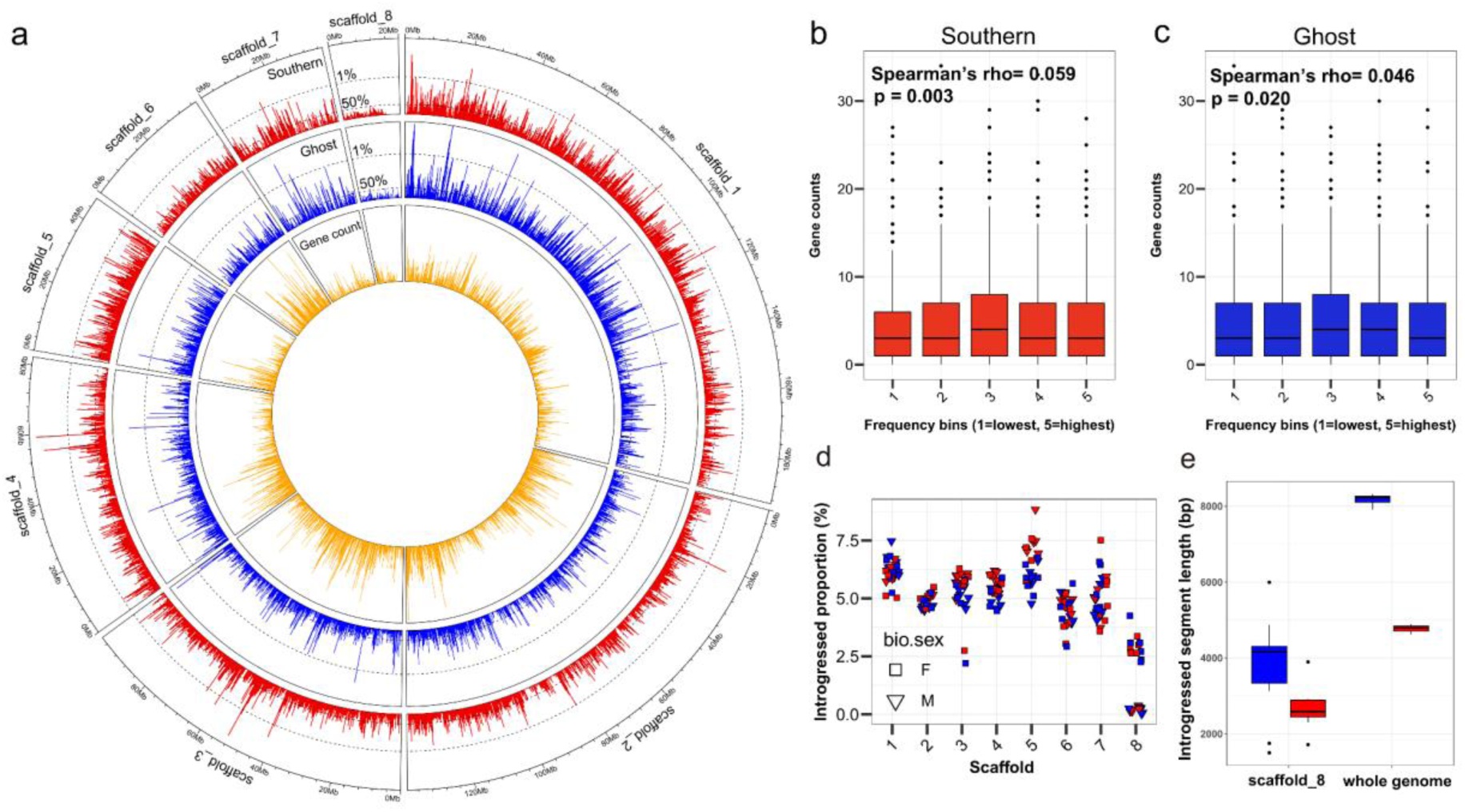
Selection pressures on introgressed segments. **a**, Genomic distribution of introgressed segments in Qingdao *P. longiforceps*. Outermost circle: Mean frequency of southern segments in each 250-kbp bin along the genome. The y-axis ranges from 0 to 0.36, with dashed lines marking the top 1% and 50% frequency thresholds. Middle circle: Same as the outermost circle, but for ghost segments. Innermost circle: Gene counts per 250-kbp bin, with the y-axis ranging from 0 to 34. **b**, Relationship between the frequency of southern segments and gene counts per 250-kbp bin. The correlation was assessed using Spearman’s rank correlation coefficient, with two-sided *P* values shown. For visualization, data were grouped into five equally sized bins based on frequency. **c**, Same as **b**, but for ghost segments. **d**, Proportion of introgressed tracts per scaffold. Each point represents a Qingdao individual, with shape indicating biological sex (female: square, male: downward triangle) and color representing origin of introgressed segments (southern: red, ghost: blue). **e**, Mean length of introgressed segments per individual in scaffold 8 and the whole genome. Colors indicate origin of introgressed segments.

**Extended Data Fig. 4.**
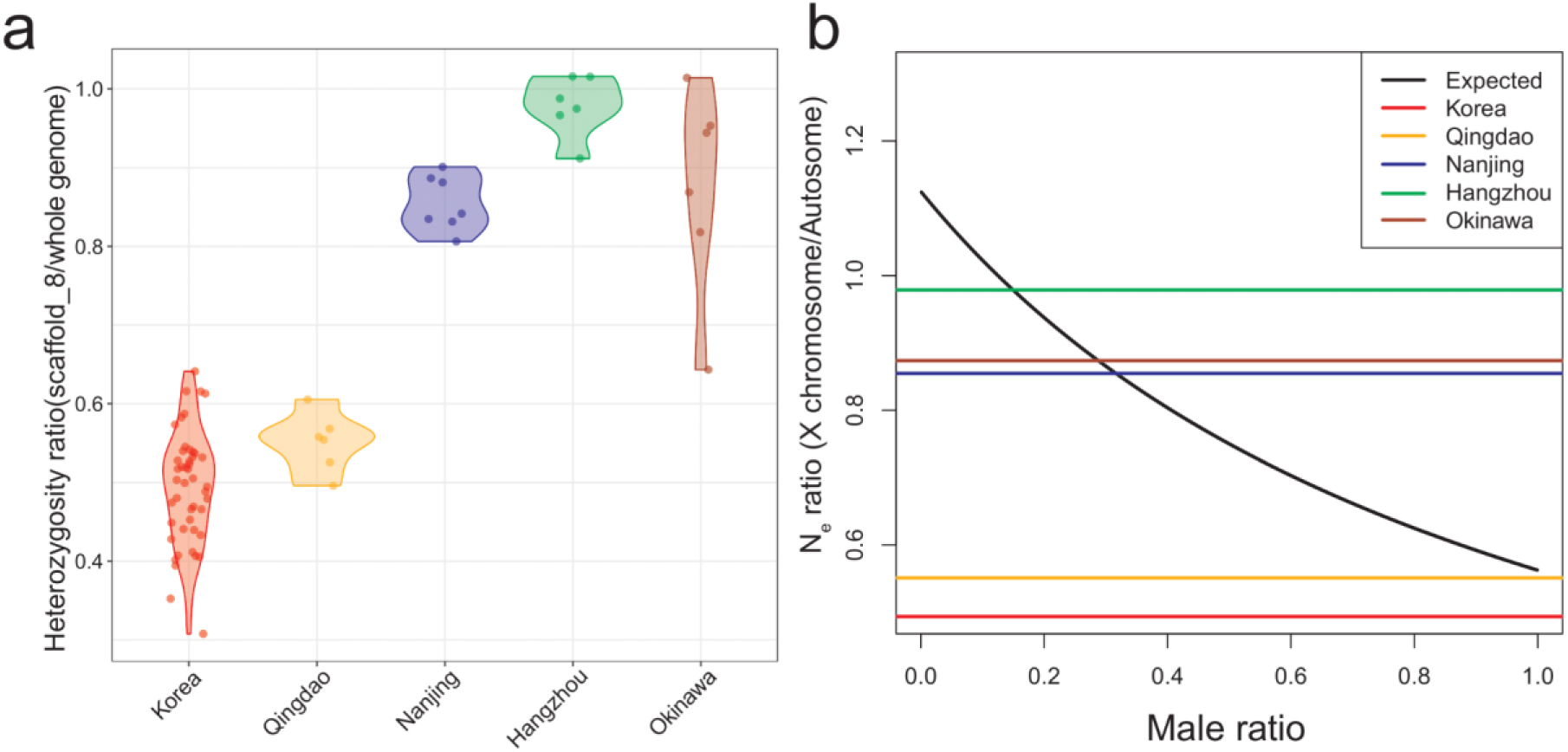
Extreme depletion of genetic diversity on scaffold 8 in northern populations. **a**, Violin plot showing the ratio of genome-wide heterozygosity between scaffold 8 and the whole genome for each female *P. longiforceps* individual across populations. Each colored point represents an individual, with the x-axis indicating populations and the y-axis representing heterozygosity ratios. **b**, Expected effective population size (N_e_) ratio between the X chromosome and autosomes under neutrality, plotted against the proportion of males contributing to reproduction (black solid line). The effective population size is calculated as: Autosome: 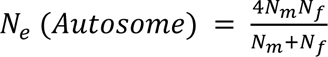. X chromosome: 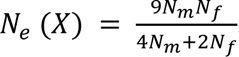, where N_m_ and N_f_ represent the number of reproducing males and females, respectively. Colored lines indicate the observed heterozygosity ratio between scaffold 8 and the whole genome for each population.

