## Supplementary material for "A deep population stratification and ongoing local adaptations of the range-expanding lovebug *Plecia longiforceps*": Fig. S

**This PDF file includes:**

Supplementary Figures 1 to 10

**Other Supplementary Materials for this manuscript include the following:**

Supplementary Tables 1 to 9

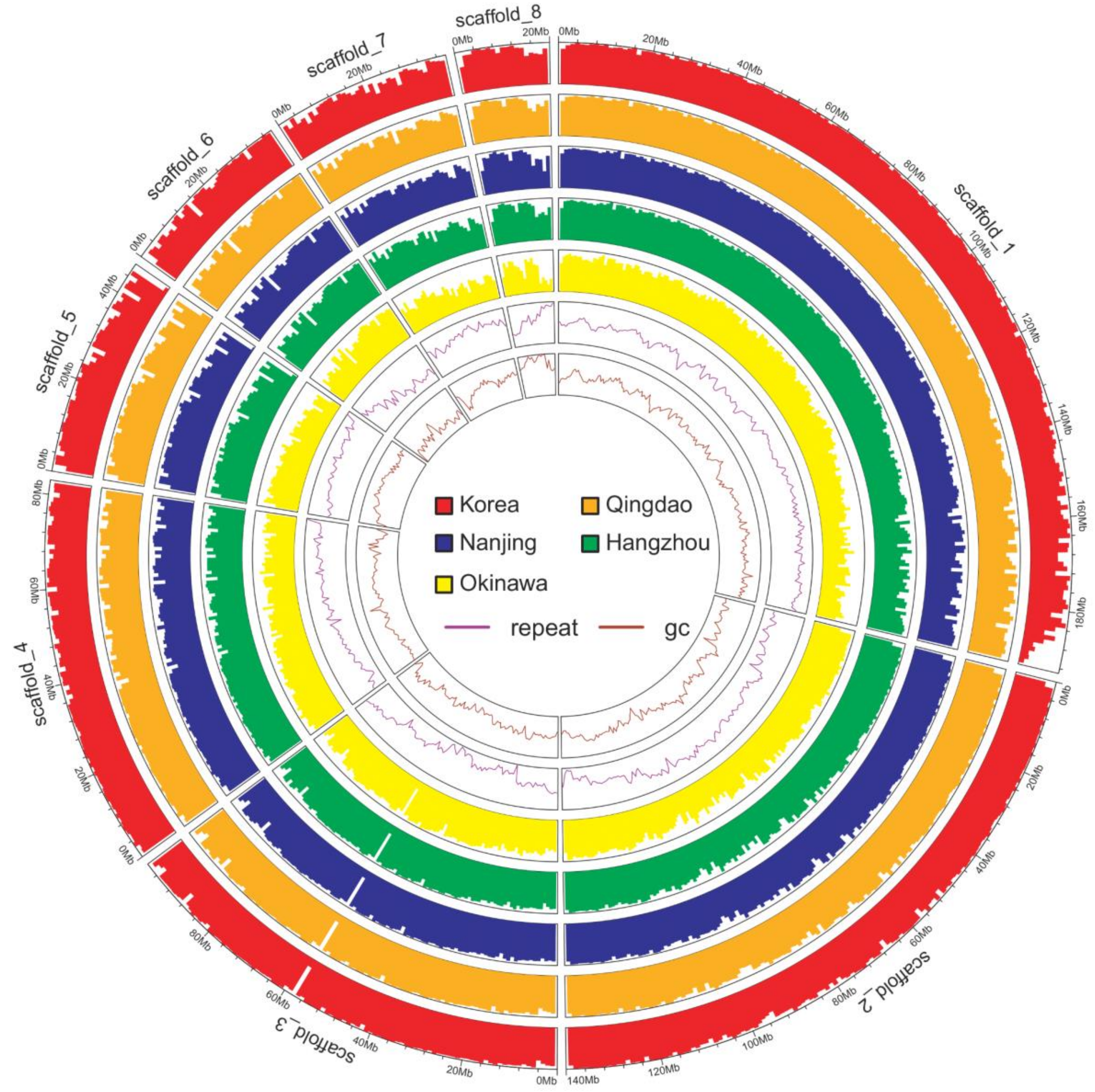


**Figure S1. The genome-wide distribution of mappability, repeat and GC contents of *Plecia longiforceps*.** The proportion of mappable 1 kb windows, repeat content, and GC content in 1Mb-bin is reported along the *P. longiforceps* genome. Five outer circles show the proportion of mappable 1 kb windows for each population—Korea, Qingdao, Nanjing, Hangzhou, and Okinawa—represented by different colors. The y-axis ranges from 0 to 1. Two inner circles display repeat content (purple line) and GC content (brown line). The y-axis for repeat content is scaled from 0.34 (minimum) to 1.00 (maximum), while the y-axis for GC content ranges from 0.26 to 0.41. Inset provides the legend for population colors, repeat content, and GC content. A mappable 1 kb window is defined for each population as a 1 kb window where the mean sequencing depth exceeds 1/5 of the genome-wide average.

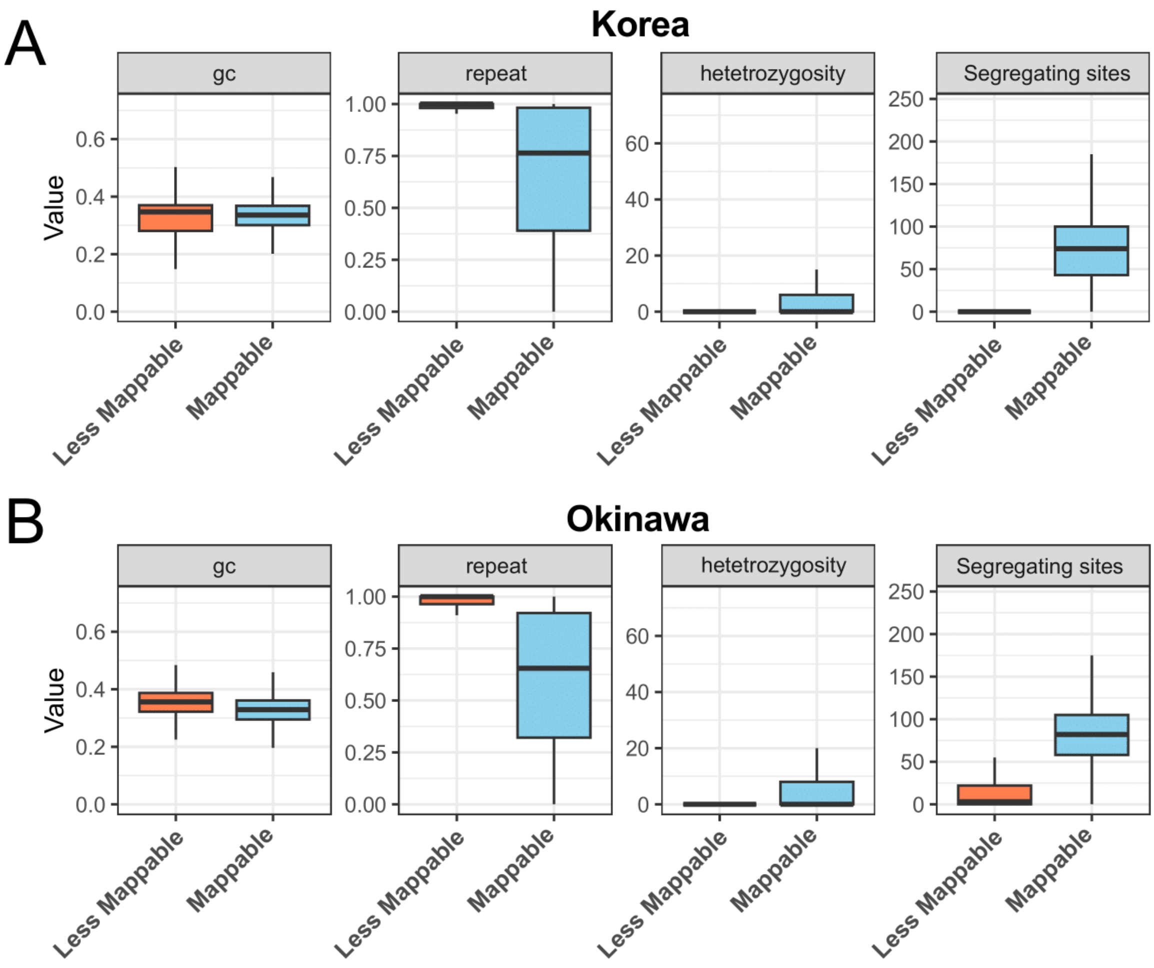


**Figure S2. Genomic properties of mappable regions. (A)** GC content, repeat content, heterozygosity, and the number of segregating sites in less mappable and mappable regions of the Korea population. A mappable region is defined as a 1 kb window where the mean sequencing depth exceeds 1/5 of the genome-wide average. **(B)** Same as (A), but for the Okinawa population. In each box plot, the box represents the interquartile range (IQR) (the 25th and 75th quartiles), and the horizon line within the box represents the median. Black-filled and open circles represent outliers (1.5 times beyond the IQR) and extreme outliers (3 times beyond the IQR), respectively.

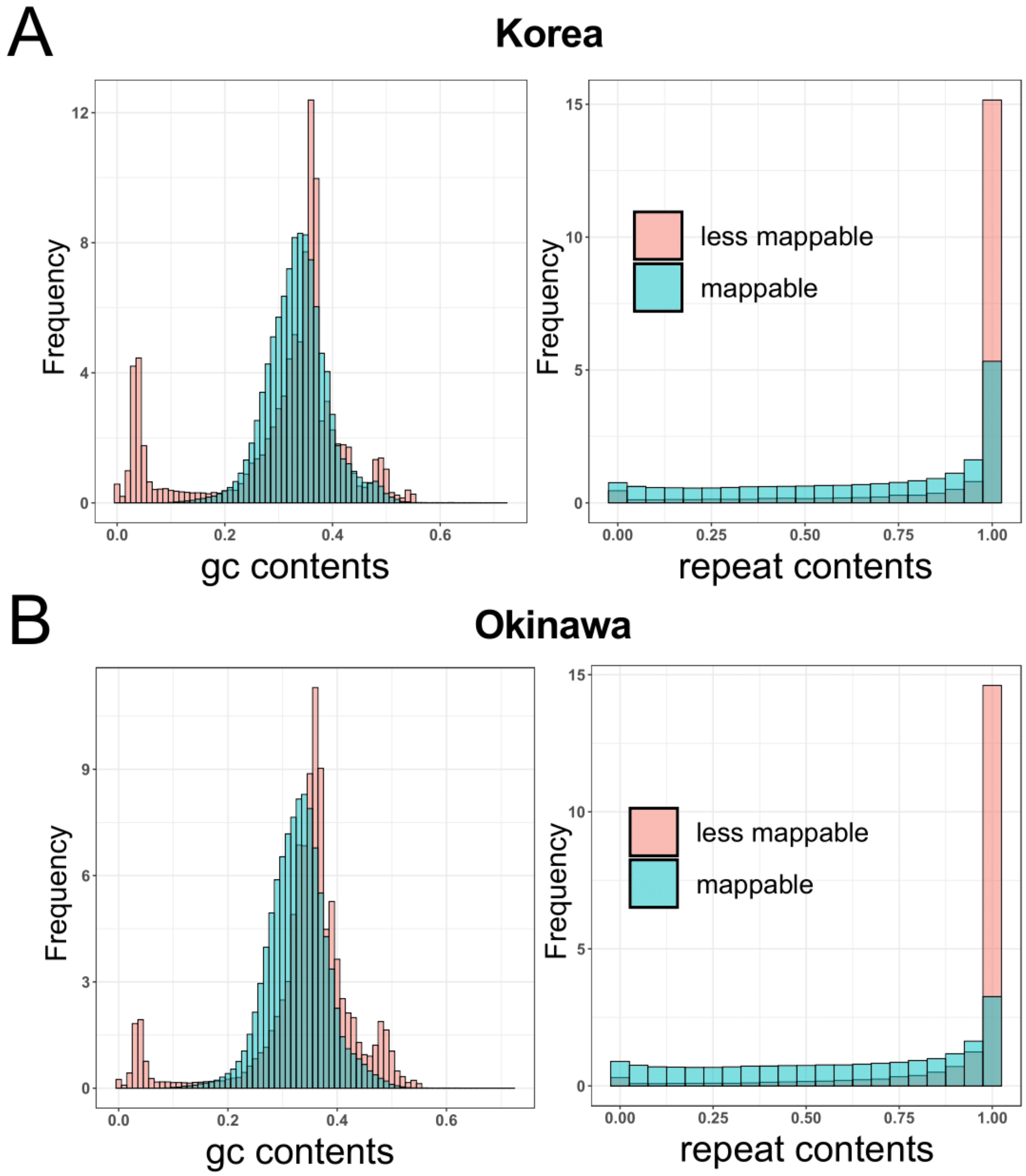


**Figure S3. Distribution of GC and repeat contents in mappable regions. (A)** GC and repeat content distribution in mappable and less mappable regions of the Korean population. Mappable regions are defined as 1 kb windows where the mean sequencing depth exceeds one-fifth of the genome-wide average. Less mappable regions are shown in transparent red, while mappable regions are marked in blue. **(B)** Same as (A), but for the Okinawa population.

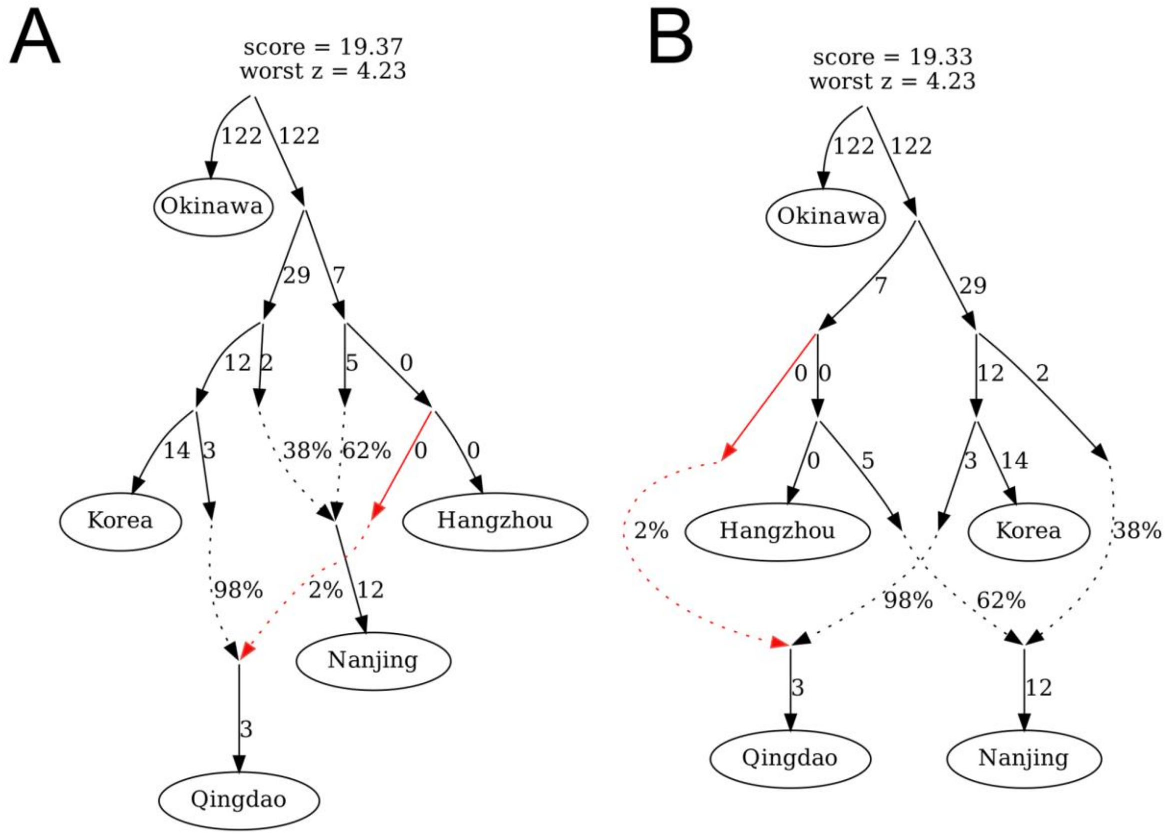


**Figure S4. Alternative population graphs of *Plecia longiforceps*.** Two alternative population graphs inferred using qpGraph depict a gene flow event from the southern lineage, unrelated to Nanjing, into the Qingdao population. In **(A),** the southern ancestry of Qingdao is closer to Hangzhou compared to that of Nanjing. In **(B),** the southern ancestry of Qingdao serves as a common outgroup to both Hangzhou and the southern ancestry of Nanjing. In both panels, the southern lineage contributing to Qingdao is marked as red. Edge lengths correspond to *F*st × 1000. The graph's log-likelihood score and the largest residual deviation between observed and expected *f*-statistics ("worst *z*") are shown above the graph.

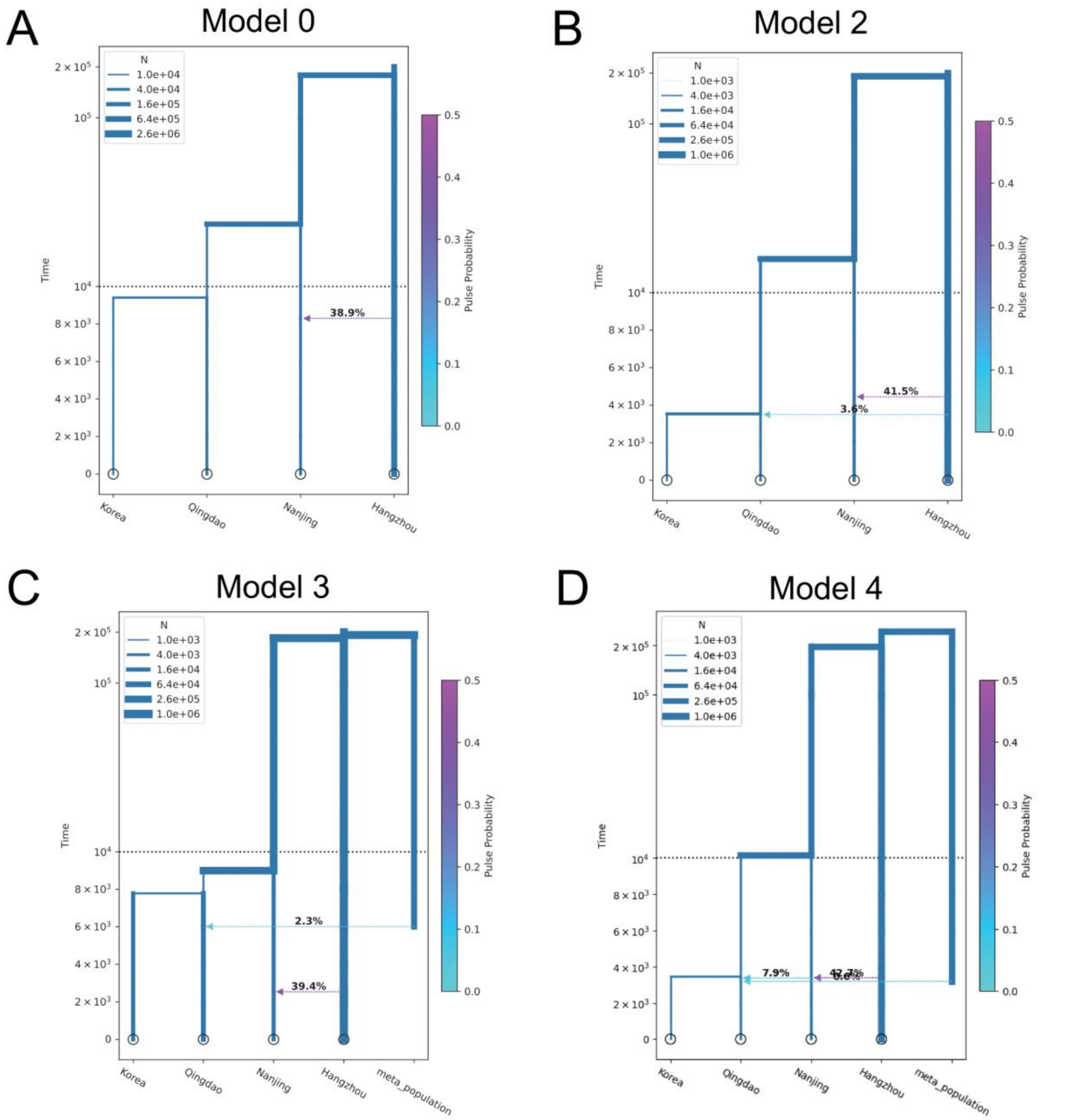


**Figure S5. Alternative demographic models inferred using Momi2. (A)** Null model representing Qingdao as a sister clade to Korea. **(B)** Model with gene flow from Hangzhou into Qingdao. **(C)** Model with gene flow from a meta-population, diverged from all sampled lineages, into Qingdao. **(D)** Model depicting Qingdao as a three-way admixture of Korea, Nanjing, and the meta-population.

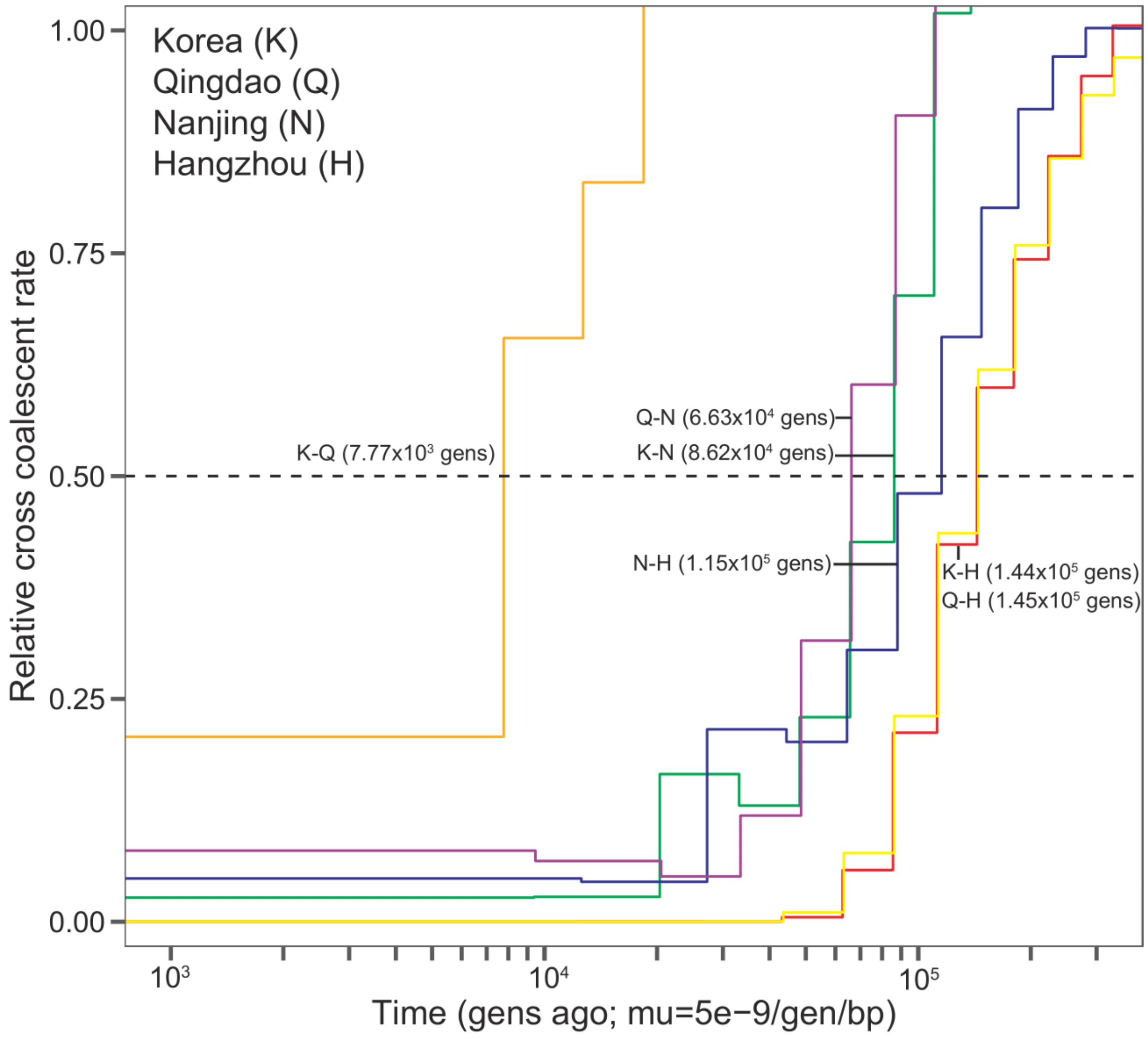


**Figure S6. Divergence time estimates based on relative cross-coalescence rates.** MSMC2 was used to infer the approximate divergence time between four populations (Korea, Qingdao, Nanjing, and Hangzhou) through cross-population analysis. The relative cross-coalescence rate (rCCR) was computed for all population pairs using: 1) The within-population coalescence rate estimated from autosomes. 2) The between-population coalescence rate estimated from scaffold 8 of males to minimize phasing errors. The divergence time between two populations was estimated as the time interval at which the rCCR reached 0.5. The coalescence rate of scaffold 8 was not scaled, as the ratio of effective population size between the whole genome and scaffold 8 varies across populations, ranging from 0.5 to 1.1.

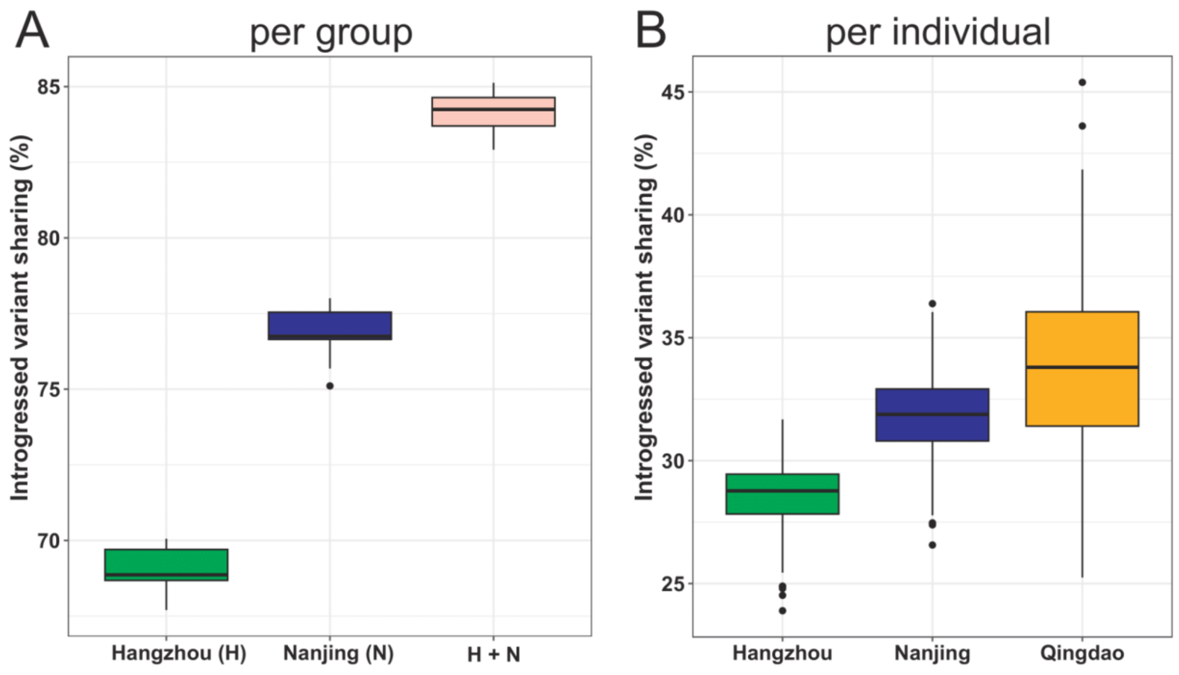


**Figure S7. Patterns of introgressed variants sharing. (A)** Proportion of introgressed variants in Qingdao individuals that are shared with at least one individual from each group: Hangzhou, Nanjing, and Hangzhou + Nanjing. **(B)** Proportion of introgressed variants in Qingdao individuals shared with each individual in Hangzhou, Nanjing, and Qingdao. Each population is represented by a different color. In each box plot, the box indicates the interquartile range (IQR) (25th–75th percentiles), with the horizontal line inside the box representing the median. Black-filled circles denote outliers (1.5× IQR), while circles indicate extreme outliers (3× IQR).

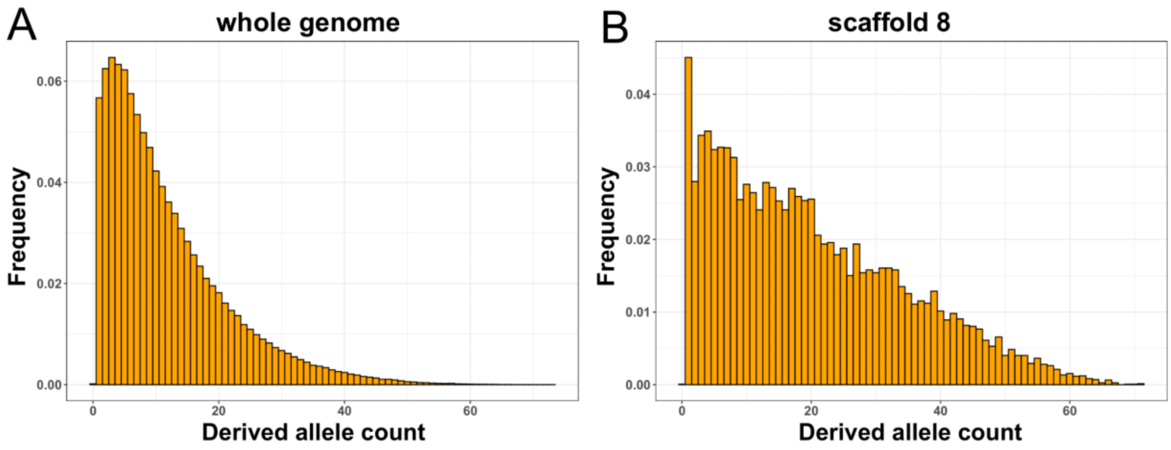


**Figure S8. Derived allele frequency spectrum of introgressed variants in the Qingdao population.** The derived allele frequency spectrum of introgressed variants identified in Qingdao individuals is shown for the whole genome (**A**) and scaffold 8 (**B**). The spectrum is calculated for southern populations (Nanjing and Hangzhou) and Qingdao.

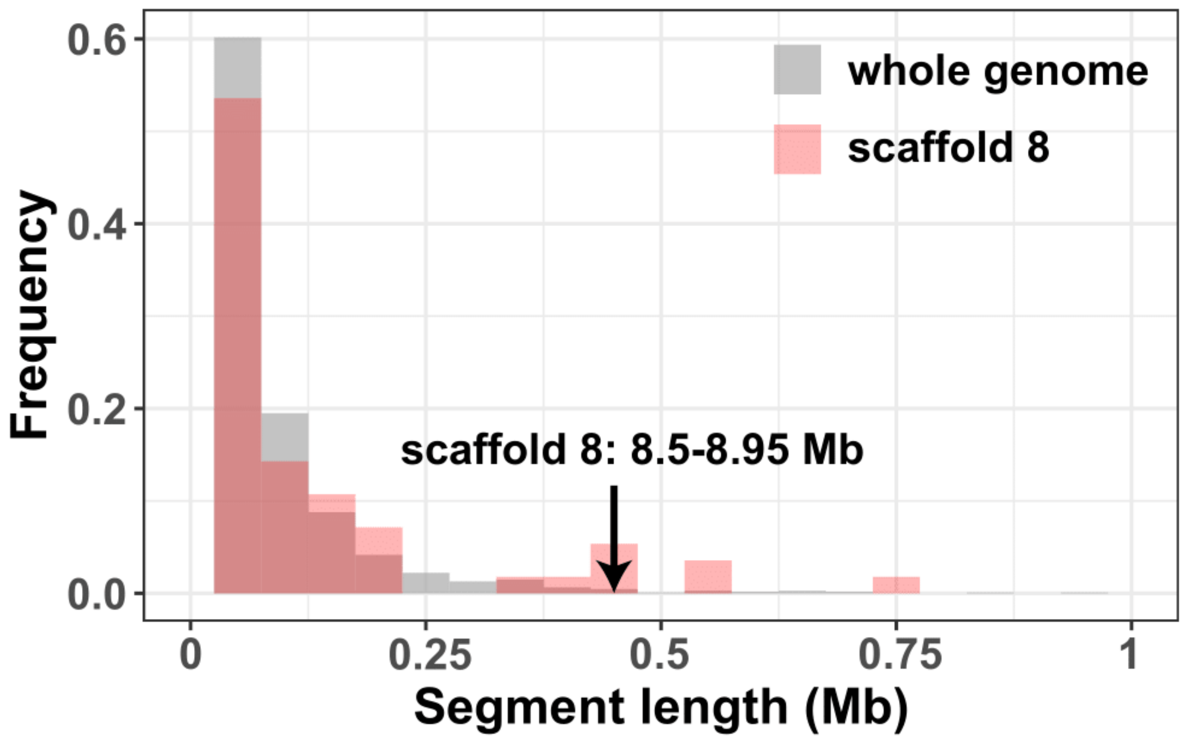


**Figure S9. Histogram of introgression desert lengths.** The histogram illustrates the length distribution of introgression deserts, representing genomic regions depleted of introgressed segments (frequency < 0.005) in Qingdao individuals. The frequency of introgressed segments was calculated using 50 kb non-overlapping windows. For clarity, we excluded a long introgressed desert on scaffold 5 (45 –46.9 Mbp). A desert on scaffold 8 (8.5–8.95 Mbp), containing windows with the strongest selective signal in northern populations (Korea and Qingdao), is marked with a directional arrow.

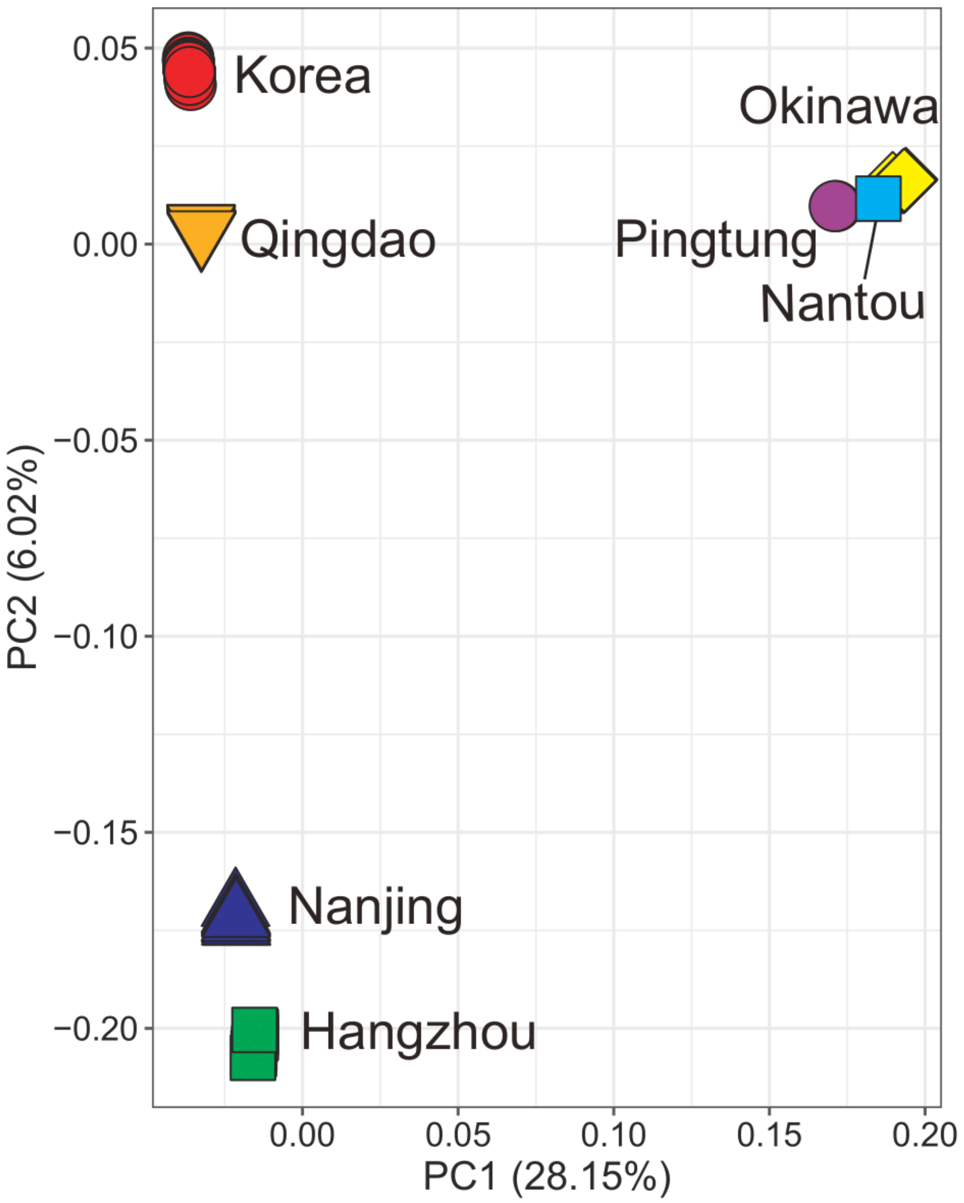


**Figure S10. Principal component analysis (PCA) of 150 East Asian *P. longiforceps* pseudo-haploid genotypes.** Each individual is represented by a color-filled symbol corresponding to its group. The proportion of variance explained by each principal component is indicated on the axis labels.
